# Single-cell analysis of EphA clustering phenotypes to probe cancer cell heterogeneity

**DOI:** 10.1101/635102

**Authors:** Andrea Ravasio, Myint Zu Myaing, Shumei Chia, Aditya Arora, Aneesh Sathe, Elaine Yiqun Cao, Cristina Bertocchi, Ankur Sharma, Bakya Arasi, Vin Yee Chung, Adrianne C. Green, Tuan Zea Tan, Zhongwen Chen, Hui Ting Ong, N. Gopalakrishna Iyer, Ruby YunJu Huang, Ramanuj DasGupta, Jay T. Groves, Virgile Viasnoff

## Abstract

Eph receptors, a family of receptor tyrosine kinases, play a crutial role in the assembly and maintenance of healthy tissues. Dysfunction in Eph signaling are causally and correlatively associated with cancer progression. In breast cancer cells, dysregulated Eph signaling has been largely linked to alterations in receptor clustering abilities. In the present study, we implemented a single-cell assay and a scoring scheme to systematically probe the spatial organization of activated EphA receptor in carcinoma cells of different origin. Using this assay, we found that cancer cells retained EphA clustering phenotype upon cell division for several generations and degree of clustering reported for population as well as single-cell migration potential. Finally, using patient-derived cancer cell lines, we probed the evolution of EphA signalling in cancer cell populations that underwent metastatic transformation and acquisition of drug resistance. Taken together, our simple and scalable approach provides a reliable quantitation of EphA associated gene expression and phenotypes in multiple carcinomas and can assay the heterogeneity of cancer cell populations in a cost- and time-effective manner.

## Introduction

Receptor Tyrosine Kinases (RTK) are critical for tissue homeostasis [1–3] and their dysfunction is associated with cancer progression [4,5]. Eph receptors, which comprise the largest family of receptor tyrosine kinases, interact with ephrin ligands expressed on the surfaces of neighboring cells. In particular, an increase in EphA activity has been implicated in many cancer types [6] including 40% of breast cancers [7,8]. This juxtacrine receptor-ligand signaling creates significant opportunities for spatial and mechanical constraints at the cell-cell interface to become intermingled with signaling behavior [9–11]. Recent studies on breast cancer cell lines reported that cytoskeleton remodeling and EphA2 signaling interact in a feedback loop [12] to control the clustering of EphA2, which specifically binds to ephrinA1 ligands [13–16]. Alterations in this feedback loop result in changes in receptor aggregate morphologies, which, in turn, lead to phenotypic changes in cell morphologies and behaviors [17–19]. Seminal reports strongly suggest that the activities of EphA2, and of other Eph receptors, [20,21] are regulated by their ability to spatially cluster in a cytoskeleton-dependent way. The cluster morphologies correlate with the invasion potentials of breast cancer cell lines at the population level [13,15]. Hence, cluster morphology could serve as a proxy for the level of activation of downstream signaling cascades. Therefore, developing a generic scoring scheme for individual cells could provide a robust phenotypic assay to ascertain the heterogeneity of cellular functional states within a cancer cell population, which could be compared across cancers of different origin. In particular, it could probe intra-tumor functional heterogeneity of EphA signaling in a cost- and time-effective manner. Such assays could also complement single-cell transcriptomic approaches by integrating gene expression profiles with essential information about cell behavior and cell-state in an unbiased way at the single cells’ level. Furthermore, such assays could pave the way for the development of a whole toolbox for screening phenotypes associated with activities of membrane receptors. In this paper, we developed a simple and scalable phenotypic assay and established a general scoring scheme to quantify EphA clustering induced by ephrinA1 ligation that we standardize amongst a variety of human cancer cell lines of different origins. We demonstrated that cluster morphology is an inheritable trait that propagates over cell divisions for several generations, and that it correlates with the migratory potential of single cells. Finally, we validated that our assay predicts alterations in EphA activation pathway in patient derived cells (PDCs) from primary, metastatic, and drug-resistant tumor cell populations. Altogether our study underscores the utility of the EphA-based phenotypic assay as a novel tool to interrogate the evolution of cancer cell states during tumor progression in clinical samples.

## Results

### Working principles and work flow of the assay

In this assay, we used supported lipid bilayers functionalized with fluorescently-tagged (Alexa 568) ephrinA1 to allow for cytoskeleton-dependent clustering of EphA receptors. Upon ligand binding, the receptors oligomerize and get trans-phosphorylated. The mobile ligand/receptor pairs then aggregate into clusters driven by cytoskeleton rearrangements [15]. Using single-cell image analysis, we quantified cluster morphologies based on scalar scores that reflecte their degree of aggregation. The distribution of individual scores provided information about intra-tumor heterogeneity. Figure 1a summarizes the experimental scheme that is detailed in the **Methods**. In brief, a silicon gasket (20 × 20 × 0.3 mm), comprising of 9 independent recording chambers (3 × 3 mm), was layered on a clean, hydrophilic glass coverslip (**Supplementary Figure. 1a**). We generated supported bilayers (96 mol% DOPC and 4% DOGS-NTA-Ni) by standard Small Unilamellar Vesicle (SUV) deposition (**Methods**) in millimeter-sized chambers. Polyhistidine-ephrinA1 labeled with Alexa 568 were conjugated with the NTA lipid moieties at saturating densities (4%). We optimized the size of the chambers and the spatial arrangement of the duplicates to achieve high reproducibility and to allow internal quality controls (**Supplementary Figure 1b** and **c**). Following sedimentation, suspended cells (approximately 100 cells per chamber) establish contact with the bilayer within 30-60 seconds (Figure 1b). Upon establishment of contact, EphA receptors on the cell surfaces cluster and engage with the ligands freely diffusing on the bilayer (Figure 1c). For all the cell types we tested, receptor aggregates reached a steady morphology within a few minutes to half an hour. Thereafter, these structures remained largely unaltered for up to one hour. Our samples were fixed after one hour to capture statistically significant differences in EphA clustering morphologies at steady state. Individual imaging of 150-200 cells typically took an hour. In this study, a manual implementation of the assay proved sufficient. Optimizing automated imaging and analysis of larger cell populations allowed by the scalable format of the assay (Figure 1a-c) was left for further development.

**Figure 1.**
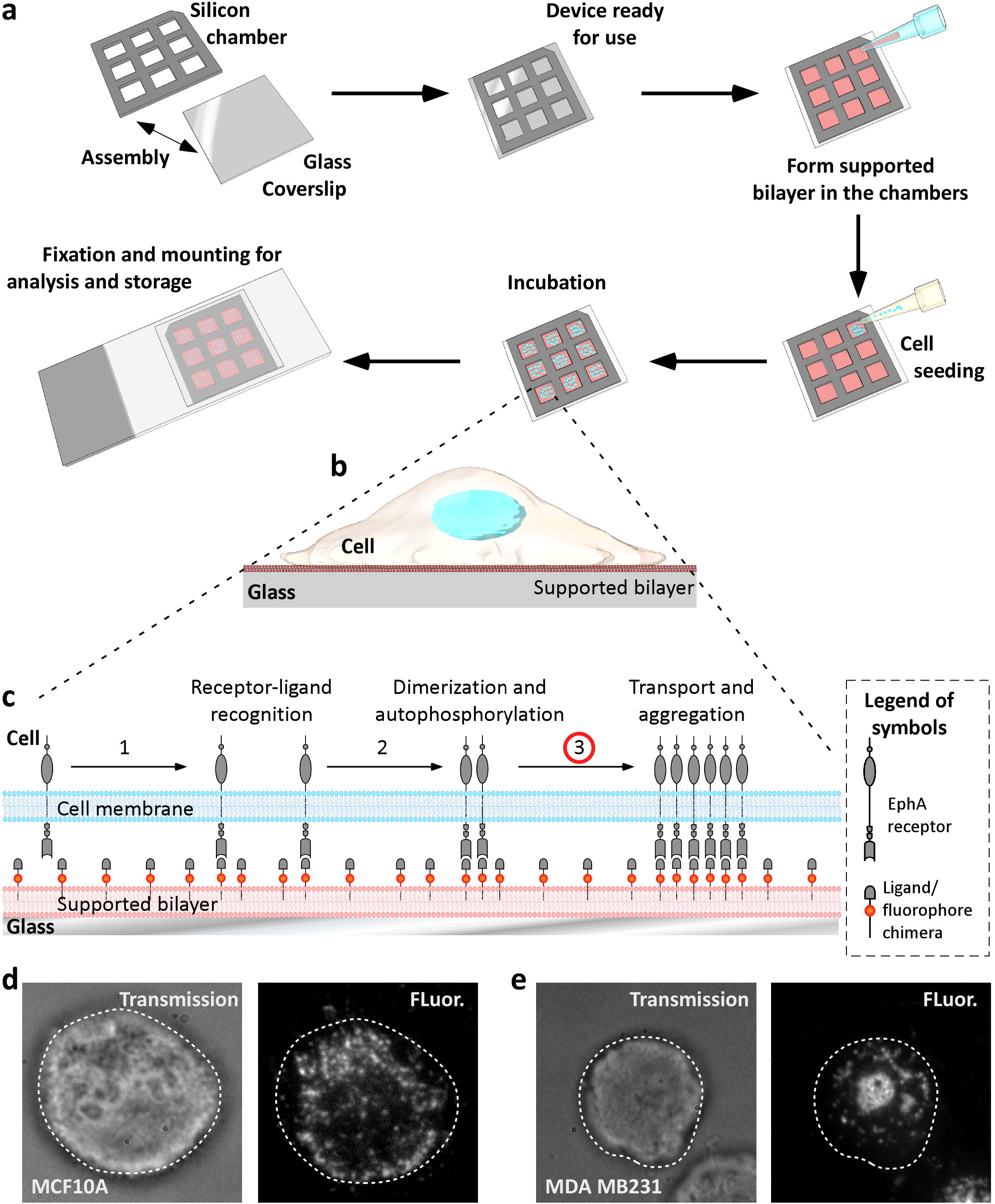
Illustration of multi-well ephrin clustering assay. **a)** Schematic representation of the experimental procedure. A silicon gasket comprising of nine chambers is placed on a clean coverslip. The coverslip is biofunctionalized with small unilamellar vesicles to form a supported lipid bilayer. The bilayer presents ephrinA1 ligands. Cells are seeded in each chamber, left for sedimentation and incubation for an hour. The system is fixed and mounted on a coverslip for subsequent imaging. **b and c)** Cartoons illustrating the EphA clustering assay. By sedimentation, cells contact the supported bilayer functionalized with fluorescent chimera of the ephrinA1 ligand. Upon receptor/ligand recognition (step 1), active receptors dimerize and phosphorylate (step 2), initiating the signaling cascade. In response to the initial signaling, active transport of receptors created signaling clusters. Their aggregation is imaged using ephrinA1 tagged with Alexa 568. **d)** Examples of images of non-invasive breast cancer cells (left, MCF10A) and an **e)** invasive breast cancer cells (right, MDA MB231) presenting different cluster morphologies. Scoring of individual cells based on quantification of the cluster morphologies is used as a proxy to define the heterogeneity of cellular states within a population.

### EphA cluster dynamics and morphologies for cells of different cancer origins

We first tested whether cells originating from different cancer types displayed variations in EphA clustering, as seen in breast cancer cell lines [15] (Figure 1d and e). Figure 2a displays the cluster morphologies obtained for seven ovarian (PEO1, SKOV3, OVCA420, OVCA429, A2780, HeyA8, and OVCAR10), one lung (A549), one gastric (MKN28) and three breast cancer cell lines (MCF10A, MCF7, MDA MB231). Reflection Interference Contrast Microscopy (RICM) images confirmed that the EphA clusters localized in regions where cells established intimate contact with the bilayer (Figure 2b).

**Figure 2.**
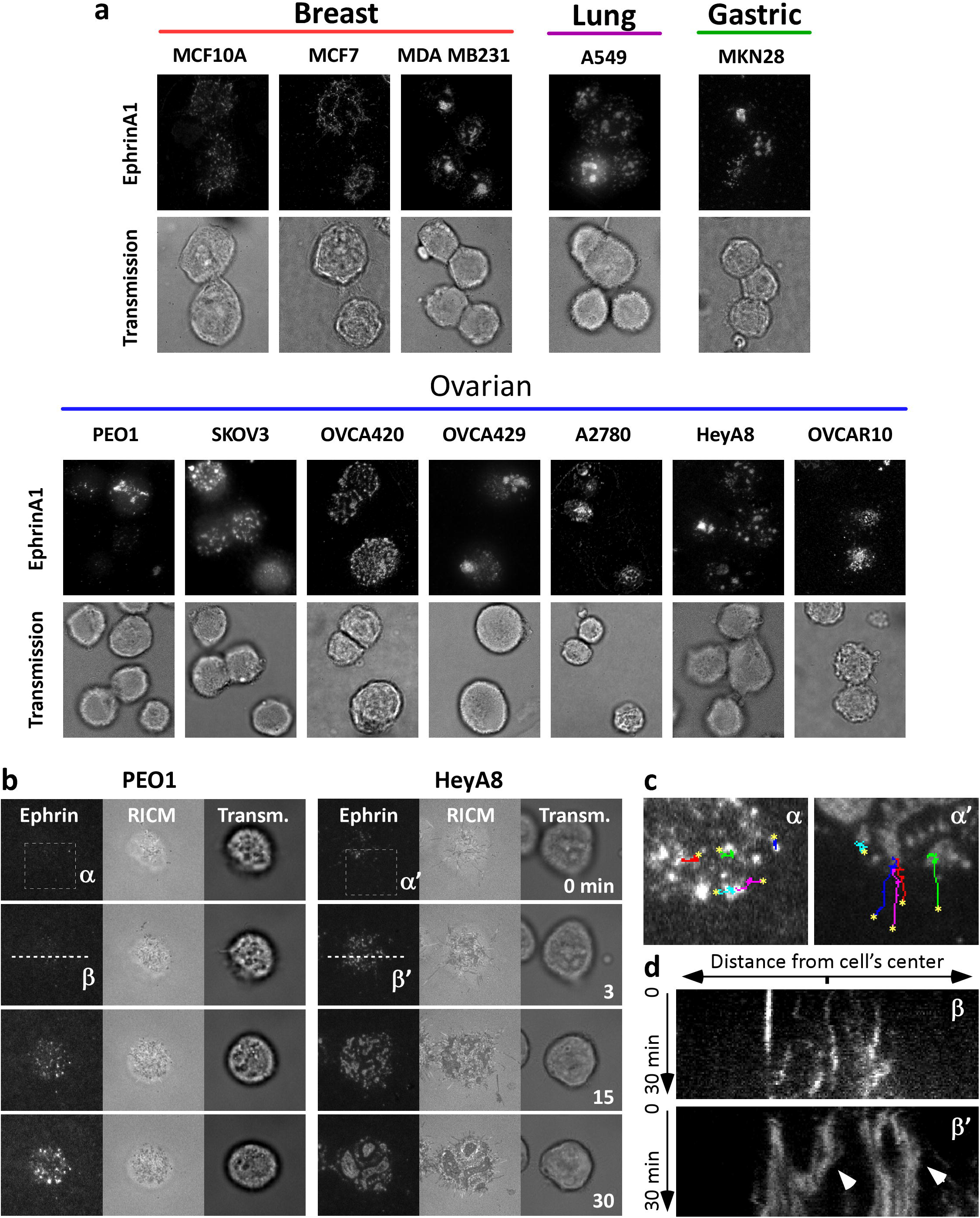
Cancer cells from various origins show morphological variability in receptor clustering. **a)** Various Eph clustering morphologies obtained across a library of cancer cells lines from various cancer origins. **b)** Time-lapse imaging of the clustering process in PEO1 and HeyA8 cells: non-invasive and epithelial-like cancer cells (PEO1) showed limited ability to form clusters as compared to the more invasive and mesenchymal-like cancer cells (HeyA8). **c)** Enlarged view from boxes (α) and (α’) in b. Trajectories of fluorescent puncta are shown by the colored lines. Initial location of the puncta is indicated by an asterisk. **d)** Time evolution (kymograph) of fluorescence along dashed lines (β) and (β’) in b. Whereas Eph clusters remained scattered in PEO1, further aggregation and merging of large clusters is observed in HeyA8 cells (arrow heads).

Interestingly, the steady state variations in cluster morphologies between cell types resulted from differences in receptor-ligand radial transport (Figure. 2b-d and **Supplementary Movies 1** and **2**). Some cell types displayed small aggregates with very little cell-to-cell variability whereas other cell types presented larger and non-homogeneous cluster morphologies. For example, PEO1 cells presented small EphA clusters with limited mobility over 30 min of observation (Figure 2c, left and kymograph in Figure 2d, top). In contrast, EphA puncta in HeyA8 cells were quickly transported towards the inner regions of the cells (Figure 2c, right), where they fused into aggregates. These aggregates further merged with other aggregates (kymograph in Figure 2d, bottom) to form a series of larger clusters (Figure 2b).

### Scoring cluster morphologies

We hypothesized that some morphological features of the EphA clusters could reflect a generic cell response to ephrinA1 binding that was independent of cancer type, whereas a different set of morphological features would be cancer-type specific. We thus probed 75 morphological descriptors of the cluster patterns for their specificity, using a combination of t-distributed stochastic neighbor embedding (t-SNE) algorithm and a machine-learning method (Random Forests) (see **Supplementary Material, Supplementary Table 1** and **2** and **Supplementary Figure 2** and **3**). This unbiased study singled out the intensive descriptor 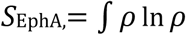, where 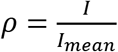, is the ratio of the fluorescence intensity value *I* of each pixel divided by the average fluorescence intensity *I*_*mean*_ under each cell (**Supplementary Figure 4**). *S*_EphA_ emerged as the best parameter to score the scattering degree of EphA clusters while ignoring their specific spatial arrangements that mostly reflected the cancer cell type of origin (**Supplementary Figure 4** for justifications). Morphologies with scattered small puncta resulted in low *S*_EphA_ scores while morphologies with large aggregates resulted in higher scores. Figure 3a shows the distribution of single-cell scores obtained for the epithelial-type PEO1 carcinoma cells and the mesenchymal-type HeyA8 cells. We linked individual scores to the corresponding cluster images. Whereas PEO1 cells showed homogenously limited clustering, HeyA8 cells displayed a high variability in cluster sizes, which correlated with high variability in *S*_EphA_ scores. In this heterogeneous population, cells with scattered small clusters obtained low S^EphA^ values (consistently with PEO1) and cells with large aggregated clusters obtained high S_EphA_ values.

**Figure 3.**
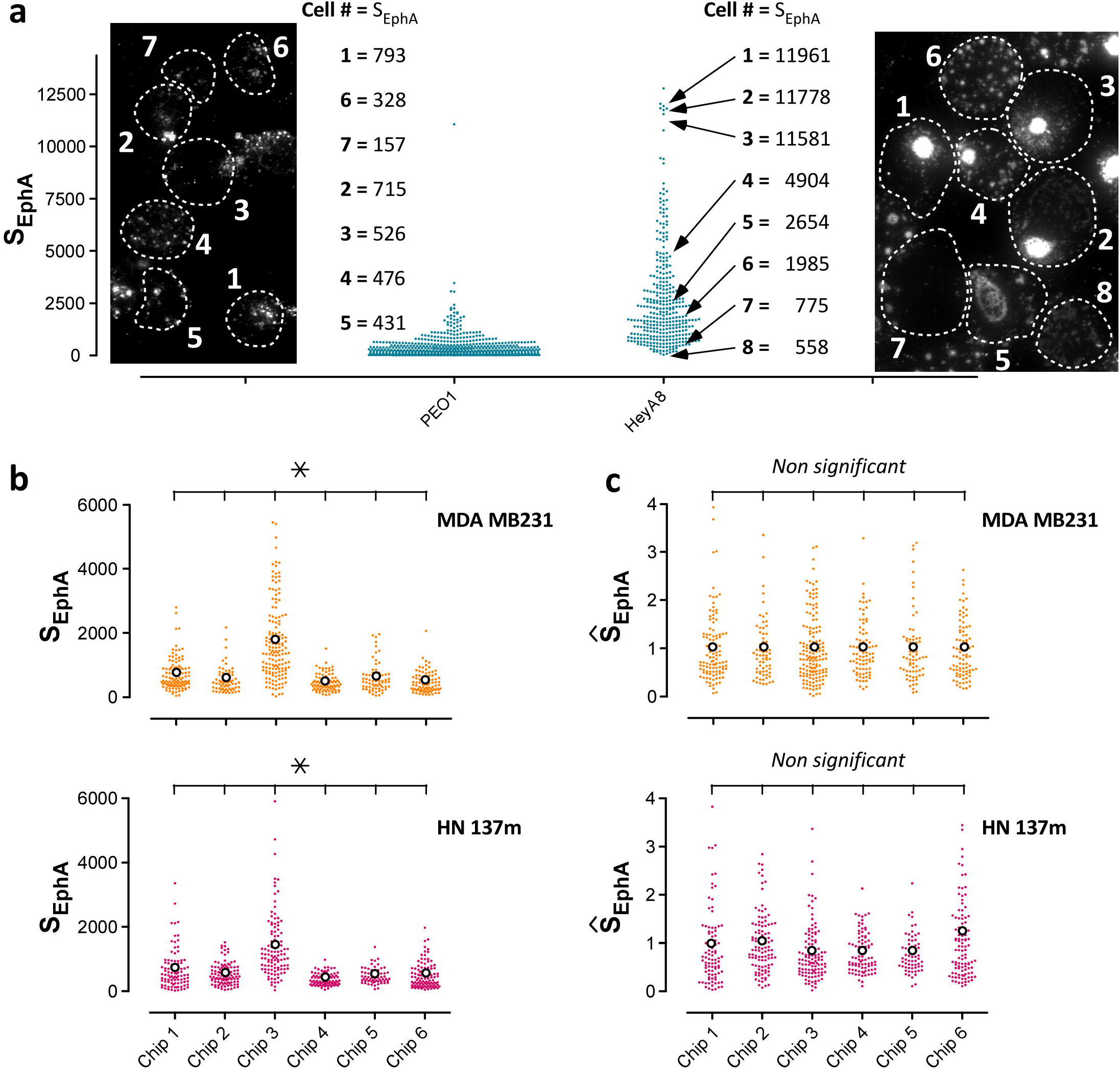
Reproducible measurement of S_EphA_ scores. **a)** Distributions of S_EphA_ scores for PEO1 and HeyA8 lines. Every PEO1 cell displays very scattered puncta. It is reflected by the low average S_EphA_ score for the population and a distribution with a small spread. HeyA8 population presents a large intercellular variability in clustering morphologies that is reflected in the high population score and a distribution with a large spread. The matching between clustering morphologies and S_EpHA_ scores is exemplified for PEO1 (7 cells) and HeyA8 (8 cells) lines. **b)** The *inter*-chip (6 chips) variability of the score distribution for MDA MB231 (top) and HN137m (bottom) cells shows significant differences between replicates. **c)** Normalization of all distributions on individual chips by the average scores of triplicates of the reference cell line MDA MB231 abolishes significant inter-chip variability. It allows quantitative comparison between chips and cell types. Each distribution is based on N>200 single-cell analysis. We established significance using the two-sample Kolmogorov-Smirnov test that probes for changes in the shape of the distribution.

### Quantitative assessment of S_EphA_ distribution for a cell population

We first tested the reproducibility of the determination of the S_EphA_ score distribution over several cell types. Repeated measurements for the same cell population on different chips resulted in qualitatively identical salient features of the distributions (narrow *vs* spread distribution). However, inter-chip variability limited the reproducibility of quantitative data. We noticed that with every chip, the experimental variations for all the tested cell lines were correlated. This suggested that quantitative changes arose from day-to-day fluctuations in the detection system rather than from cell-intrinsic noise. We alleviated these variations by implementing an intrinsic normalization scheme. On every chip, we measured triplicates of MDA MB231cells that were used as a reference cell line (**Supplementary Figure 1b**). **Supplementary Figure 1c** shows an example of the intra-chip and inter-chip variability for each triplicate in 3 independent chips. As a quality control measure for device preparation, we considered only those individual chips for which the MDA MB231 triplicates showed no significant differences (p>0.05 using a Kruskal-Wallis non-parametric test with Dunn’s multiple comparison applicable to non-Gaussian distributions). When this criterion was not met (in less than 20% of chips), the entire chip was excluded from analysis. For each selected chip, we then combined the distribution obtained for the MDA MB231 triplicates into a single distribution and used its mean value to normalize all the distributions measured on the entire chip. Figure 3b illustrates the combined distribution of triplicates of MDA MB231 cells (breast cancer) and HN137m cells (patient derived squamous head-and-neck cancer cell) run over 6 different chips. Pairwise comparison demonstrates the *inter*-chip variability observed experimentally. Due to the proportionality of the pairwise *inter*-chip variability, we could normalize all the scores for a given chip by the averaged value of the MDA MB231 distribution (N>150 cells). Normalized values are represented as 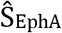, where 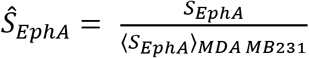. The use of a reference cell line is a simple strategy that provides dual advantage: a method for controlling chip quality and a scheme to reduce inter-chip variability below 5 % (Figure 3c).

### Inter-cell heterogeneity of clustering for cell populations from various cancer types

We then used the above approach to quantify the distribution of single-cell 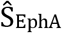 scores for the 12 cancer cell lines of breast, ovarian, lung and gastric origins (Figure 4a). Independent of the cancer type, epithelial-like cell displayed lower, narrowly-distributed 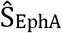 scores, whereas more mesenchymal-like cell showed higher, widely-distributed 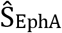 scores. Independent of their mean values, the shape of the distributions differed between the cell lines (Figure 4b). The combinatorial statistical differences between different populations are presented in **Supplementary Table 3**. While PEO1 cells predominantly displayed very low scores with a relatively long tail towards higher scores, A549 cells and MDA MB231 cells mostly displayed a symmetric distribution around their average scores. Kolmogorov-Smirnov tests revealed that considering distributions shapes together with their average values provided characteristics of each cell population with 90% accuracy (**Supplementary Table 3**). Taken together, the results shown in Figure 3a and Figure 4b suggest that the shapes of 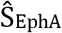 distribution, in addition to average scores, could be used to predict single-cell heterogeneity in each population, rather than being regarded as measurement noise.

**Figure 4.**
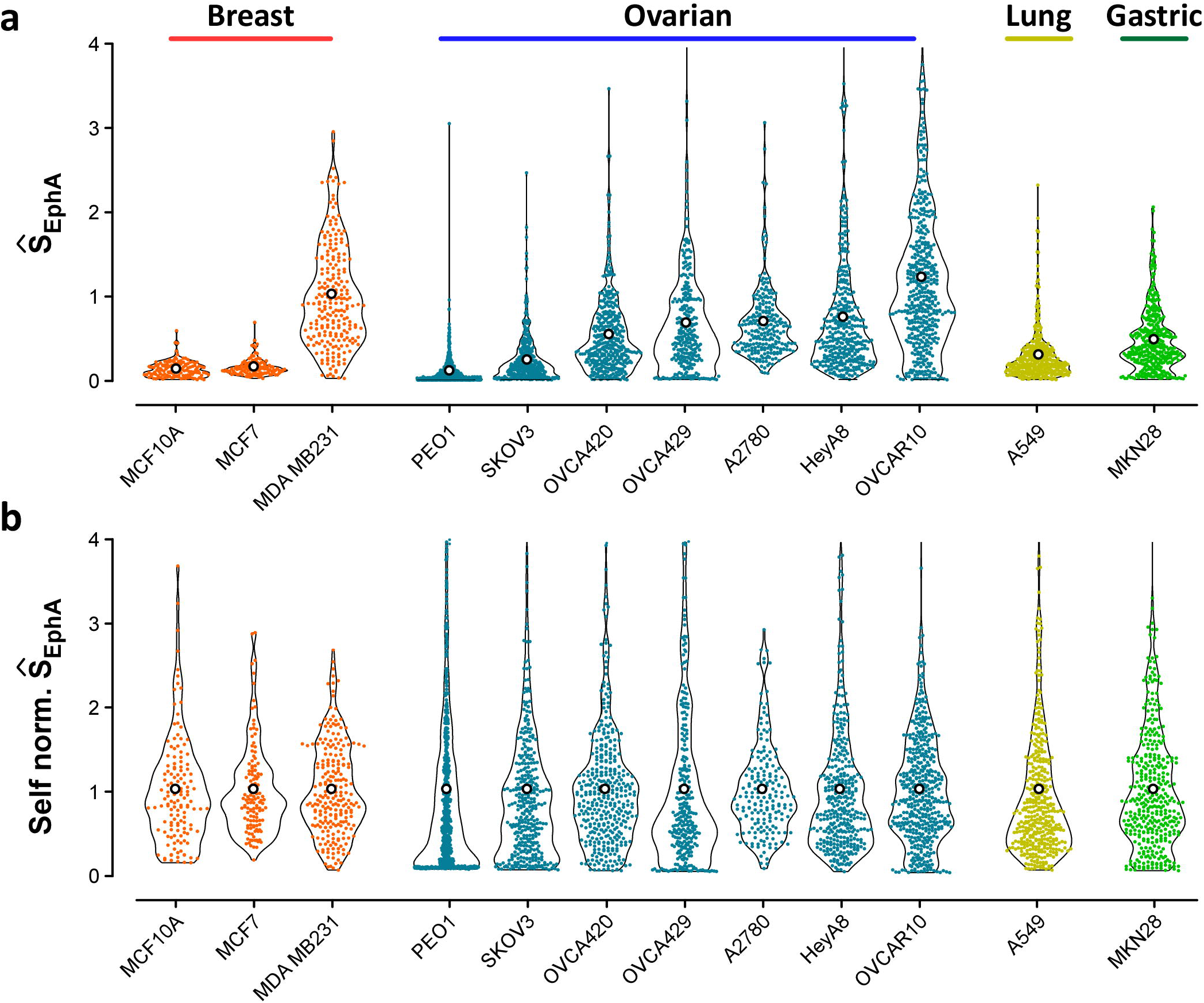
Measuring 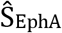 distributions across cell types. **a)** 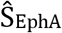 distributions for a panel of breast, ovarian, lung and gastric cancer cell lines. All cell types displayed cluster morphologies that could be quantitatively assessed in our assay, leading to significantly different average value (N>200 for each line) **b)** When the scores were normed by their own average values, they were still differently distributed around 1. For the sake of clarity, the combinatorial assessment of statistical relevance between all distributions is presented in **Supplementary Table 3**.

### EphA clustering ability is an inheritable trait

So far, we established that the distribution of EphA clustering could characterize the cell population. We then tested whether individual cell response was an intrinsic property of individual cells that could be retained even upon cell division. To this end, we adopted the following strategy. We used patient-derived primary cultures from head-and-neck tumor (specifically HN137p) [22]. We isolated and expanded 30 individual cells into clonal colonies for 10 days, following which we tested half of the cells from every colony with our assay. The other half was left to expand for another 10 days, and cells from every colony were tested again after 20 days. We then compared the distribution of scores for the original population (before cell isolation) with the distribution after 10 and 20 days for each clonal expansion. Nine colonies survived this screening process (Figure 5a).

**Figure 5.**
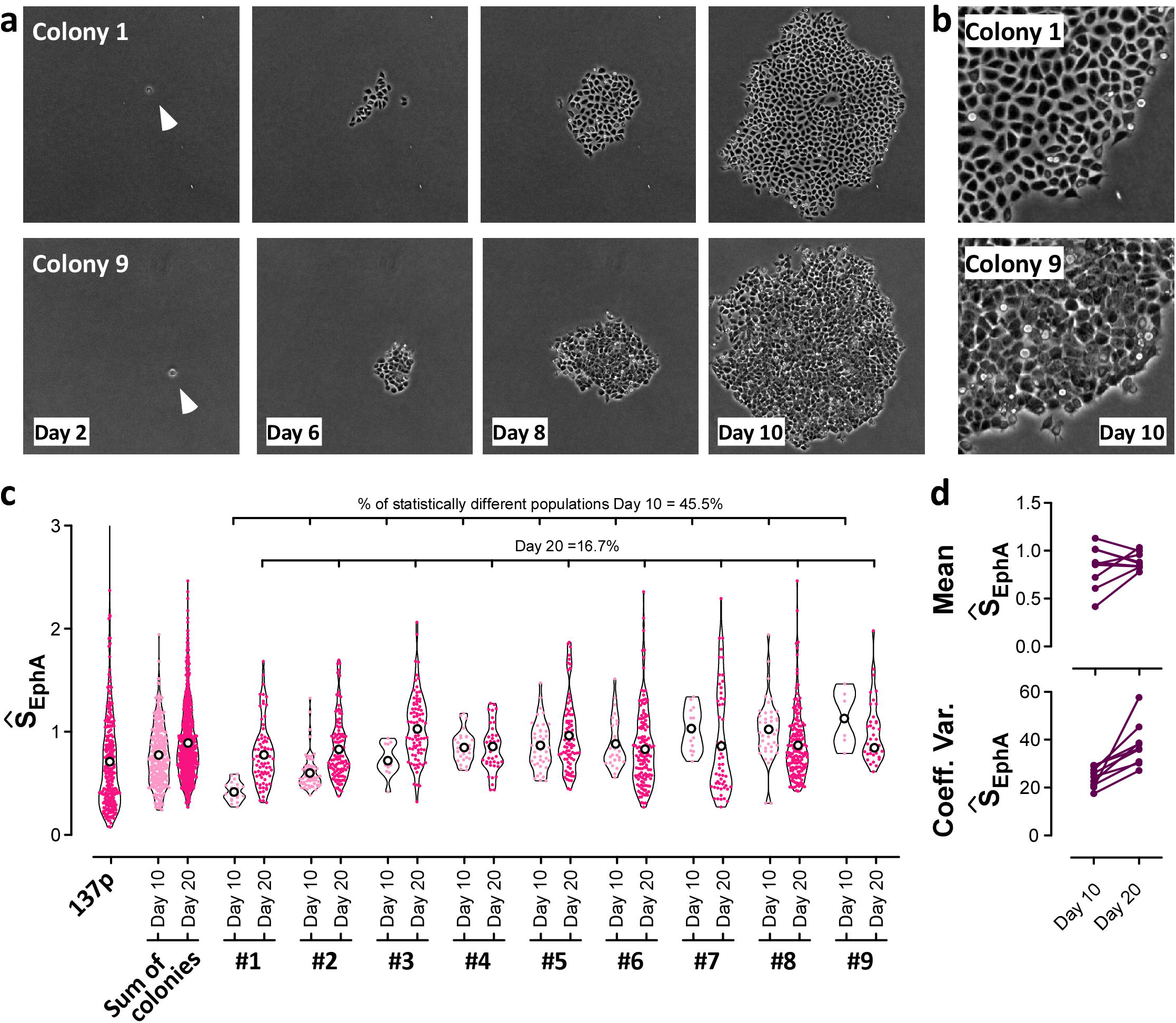
The clonal expansion of HN137p colonies demonstrates that EphA clustering is an inherited trait passed over many progenies. 30 individual cells were picked from the unsorted population of HN137p cells, out of which 9 clonal colonies could grow for 20 days **a).** Phase contrast images of the clonal colonies over time. Some colonies display a very homogeneous aspect (e.g. colony 1), some others show a noticeable spread in cell morphologies (e.g. colony 9). **b)** Close-up of colonies 1 and 9 after 20 days. **c**) 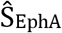 distributions for the unsorted cell population, for populations reconstituted from the 9 colonies at 10 and 20 days, and for each individual colony at both time points. Note the absence of low scores in the reconstituted populations and the significant shift towards larger score between 10 and 20 days. After 10 days, each colony retains a distribution well-centered around its average value, indicative of the conservation of the clustering properties upon cell proliferation. After 20 days though, all colony distributions converge towards the same distribution, indicating the genomic drift of the cellular states over time. **d)** Convergence of the mean score for every colony between day 10 and day 20. The spreads of all distributions increase simultaneously, indicating a larger variability in cell behavior. Statistical analysis of populations (Kruskal-Wallis with Dunn’s posttest) and confidence test (Bootstrapping) are discussed in **Supplementary Material**.

During the first 10 days (seven to nine cell divisions), we monitored the expansion of clonal colonies using phase contrast microscopy. Some colonies displayed very homogeneous morphologies (e.g. colony 1) with well-defined cell-cell contacts and smooth edges, a typical characteristic of epithelial-like cells. Other colonies, however, appeared much more heterogeneous (e.g. colony 9) with large variations in their individual cell phenotypes, a lower degree of contact inhibition, and more diffuse edges (Figure 5b).

Figure 5c compares the evolution of EphA scores for the unsorted population, and for the 9 colonies at 10 and 20 days. After 10 days, colonies with homogeneous epithelial morphologies (e.g. colony 1 or 2) displayed 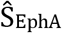 scores significantly lower than their unsorted population average. Conversely, 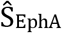 scores higher than the population average characterized the colonies with more irregular appearances (e.g. colonies 8 or 9). At day 10, each colony displayed a different average 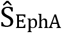 score (Figure 5d, top). However, intra-colony variability was small as shown by the narrowly-distributed scores around their respective average values (Figure 5d, bottom). After 20 days of culture, the colonies evolved towards similar 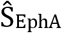 averages (Figure 5d, top), which converged towards the average value measured for the entire population. Concomitantly, intra-colony variations increased substantially as shown by the larger coefficients of variation of 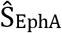 distributions (Figure 5d, bottom). The growth of clonal colonies limits the number of cells available for this analysis. However, confidence analysis (**Supplementary Material** and **Supplementary Figure 5**) demonstrates that the distribution of scores obtained for each colony cannot be interpreted (with 99.5% confidence) as a random subset of the unsorted distribution, instead they must be attributed to the phenotypes of the initial clones. Additionally, after 10 days of growth, 45.5% of the colonies significantly differed from each other (Kruskal-Wallis with Dunns post-test). This ratio dropped to 16.7% at 20 days demonstrating the “reconstitution” of the original population diversity from each clonal expansion. Of note, reconstituted populations after 10 and 20 days lacked cells with very low 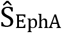 scores (below 0.2). Instead, an increasing fraction of cells presented 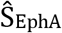 larger than 0.6. This observation was consistent with the idea that expanding clonal colonies did not favor the growth of the most epithelial-like cellular types (only 9 cells could be expanded out of the initial 30 cells, with most of them being quiescent or apoptotic). During proliferation, most epithelial phenotypes were not reconstituted, and cells appeared to drift towards the more mesenchymal cell types.

Altogether our results, suggest that EphA cluster morphology of a single cell is an inheritable trait that gradually diverges upon sequential divisions to progressively regenerate the initial population diversity.

### 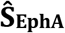 correlates with population and single-cell migration potentials

In breast cancer cells, different EphA activation levels were associated with different invasion potentials at the population level [15]. This correlation was attributed to pathways modulating the actin cytoskeleton, that could induce EphA clustering as well as promote cell migration. We therefore tested how phenotypic characterizations of EphA clustering overlapped with other standard phenotypic assay, including the migratory potential of cancer cells.

Using microchip arrays, we first measured the transcriptional levels of all EphA receptors in all our cell lines. EphA2 proved to be the most abundantly expressed receptor in eight out of the eleven cell lines (**Supplementary Table 4**). 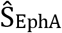 scores at the population level (**Supplementary Figure 6**) did not correlate with the mRNA levels of EphA2 and also with all EphA pulled together. The correlation greatly improved when EphA2 protein expression levels (**Supplementary Figure 6b** and **c**, Western blot) were considered. We then tested if the cellular phenotypes defined by our assay correlated with Epithelial Mesenchymal Transition scores (EMT) and their migration potential. We computed EMT scores for all cell types according to Tan et al. [23] by measuring the transcription levels of a list of 118 genes. Overall, EMT scores poorly correlated with 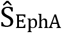(Figure 6a, left; dash line, Pearson coefficient = 0.34). However, we obtained a much higher correlation by subdividing the cell lines into two groups using 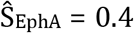 as threshold (Figure 6a, left; dotted lines, Pearson coefficients = 0.83 for group with 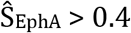, and Pearson coefficients = 0.99 for group with 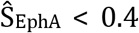). We then systematically performed wound-model experiments to assess the migration potentials of all cell lines. Interestingly, cell motility speed highly correlated with ŜEphA (Figure 6a, right; Pearson coefficient = 0.89). Furthermore, we observed that groups of cells lines with equivalent EMT scores, but different 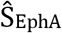 score (colored ovals in Figure 6a, left) had as a common discriminator their cell migration potential and cell lines in the group with 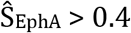 were fast and 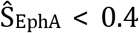 were slow migratory. These results suggest that the strong phenotypic link between EphA clustering and migration potential established for breast cancer cell lines may remain valid for cancer cell lines of different origin.

**Figure 6.**
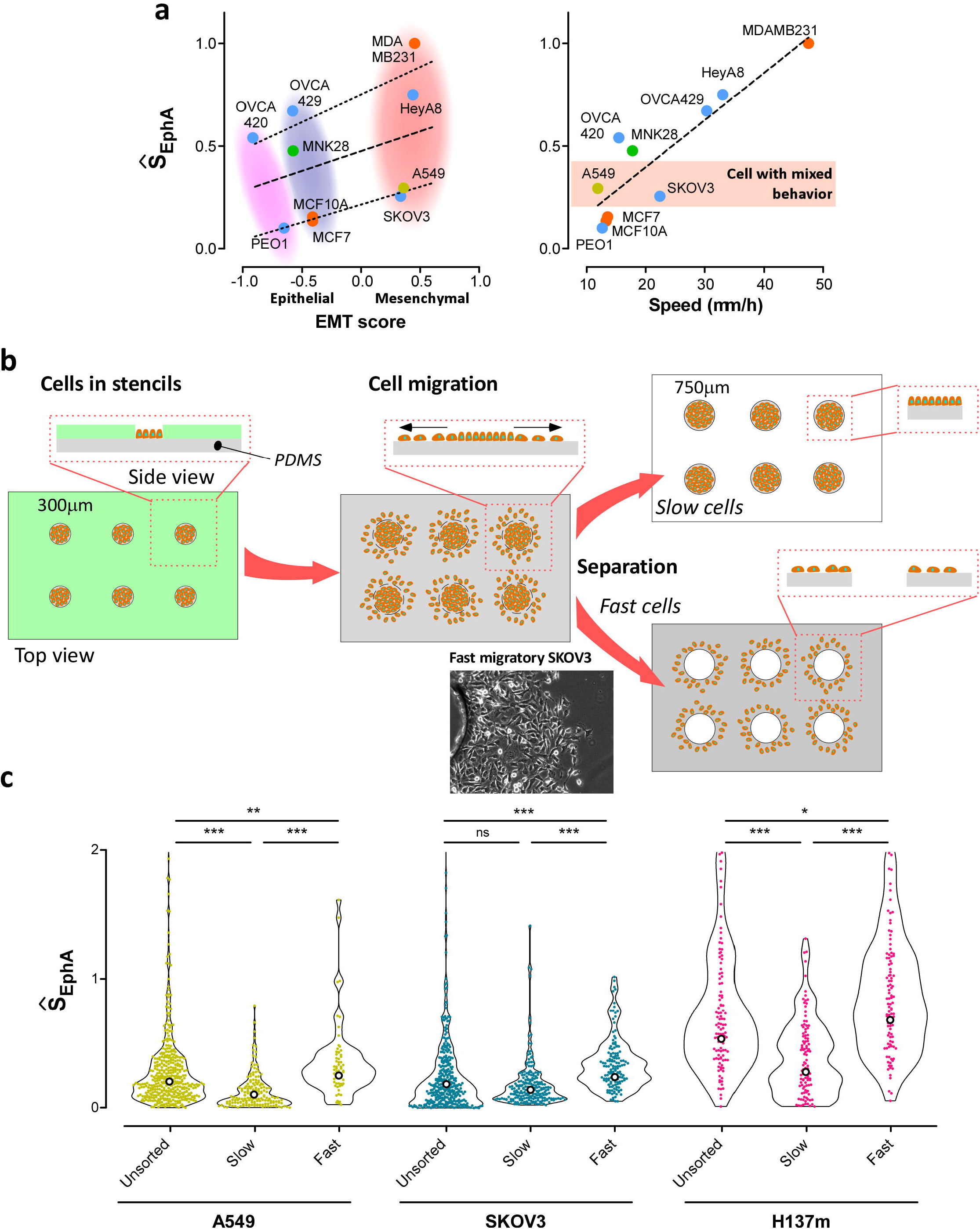
Correlation between 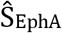 scores EMT score and migration speed. **a)** Correlation of the average 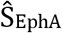 with the EMT scores (based on 118 genes). The correlation is globally poor, but it largely increases (Pearson coefficient ~0.85) if cell types are split into slow and fast migrating cells. The correlation between the average 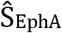 with the mean mobility of cells in the migration front of the gap-closure assay for each population is relatively high (Pearson coefficient ~ 0.89). **b)** Schematic drawing of our separation assay based on 2D migration. Cells are plated on 300 μm islands obtained by protein printing. The cells are left to migrate away from the initial island. Using a 750 μm punch, we then separated the fast migrating cells that migrated more than 750 μm away from the initial island center, from the rest. **c)** 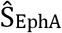 distributions for unsorted, slow and fast populations from A549 (lung), SKOV3 (ovarian), HN137m (head-and-neck) cancer cell types. Note that the fast (respectively slow) subpopulations systematically higher (respectively lower) 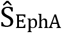 scores than the unsorted population. N>200 cells.

Some of our cell lines clearly showed heterogeneity in migratory behaviors within the cell population as it often occurs in cancer cells with intermediate EMT score (Figure 6a, right). We thus explored whether this heterogeneity was also reflected in 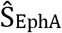 distribution patterns. In the three cell lines that we used - A549 (lung), SKOV3 (ovarian), HN137m (head-and-neck PDCs [22]) - migration speeds were substantially heterogeneous, and, correlatively, their 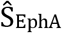 distributions were also scattered. We then used a modified wound-model assay to isolate the fast-migratory cells from the slow ones within each cell population and compared the 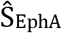 distributions between the different subpopulations (fast migratory cells, slow migratory cells, unsorted populations) (Figure 6b). We initially confined the cells on 300 micron islands on a PDMS sheet coated with fibronectin (see **Methods**) and, after removal of the confinement, we allowed migration in 2D for 3 days. We then punched-out the central part of the colonies with a 750-micron punch, thereby sorting cells that had migrated more than 250 microns away from their initial position. In all cell types, fast migrating cells displayed a 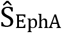 distribution with a higher average value and higher spread (Figure 6c).

These results demonstrated the correlation between EphA clustering and migration potential at the population level across diverse cancer cell types. Furthermore, enrichment of cells with high 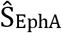 in subpopulations selected for their migration potential support the idea that 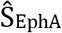 correlate with single-cell migration potential. 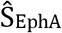 scores hence appear as a fast and reliable measurement of the migration potential of cells and heterogeneity of their invasion potential within a given population.

### Evolution of EphA signaling in cancer cell populations undergoing metastatic transformation and acquisition of drug resistance

Finally, we tested whether our assay could follow changes in the cellular phenotypes during cancer progression in patient-derived models. We have previously reported the generation of multiple primary cancer cell lines from three different head-and-neck patient tumor samples [22,24]. These include HN137, HN120 and HN148 cell lines, that were generated from the primary site of tumors (HN120p, HN137p, HN148p) or the paired lymph node metastatic sites (HN120m, HN137m, HN148m). Additionally, we also generated *in-vitro* acquired-resistance models for cisplatin for those cell lines derived from the primary site (HN137pcr, HN120pcr) [24]. Extensive characterization of these cells demonstrated that they all presented very distinct transcriptomic profiles and migratory phenotypes [22,24]. We therefore tested whether these previously reported characteristics were reflected in the distribution of 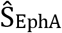 for these distinct 8 PDCs.

Figure 7a shows that for Patient HN120, the 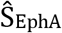 distributions from primary, metastatic and cisplatin resistant cell lines displayed increasing average values. This observation correlated with the increased migration potential observed across these cell lines [22,24]. Additionally, HN120p and HN120pcr cells displayed very different 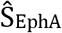 scores, corroborating the previous observation that HN120pcr cells are transcriptomically distinct from HN120p. This conclusion was also supported by comparing the 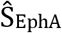 scores to the single-cell transcriptomic profiles (N>100 cells) established based on the expression levels of 71 genes, which have been reported to be downstream of EphA2 signaling cascade [15] (**Supplementary Table 5**). The transcriptional scores confirmed that all patient-derived cells lines were distinct. Importantly, EphA2 expression profile of the HN120 lines ranked similar to and correlated with the matched 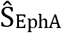 scores.

**Figure 7.**
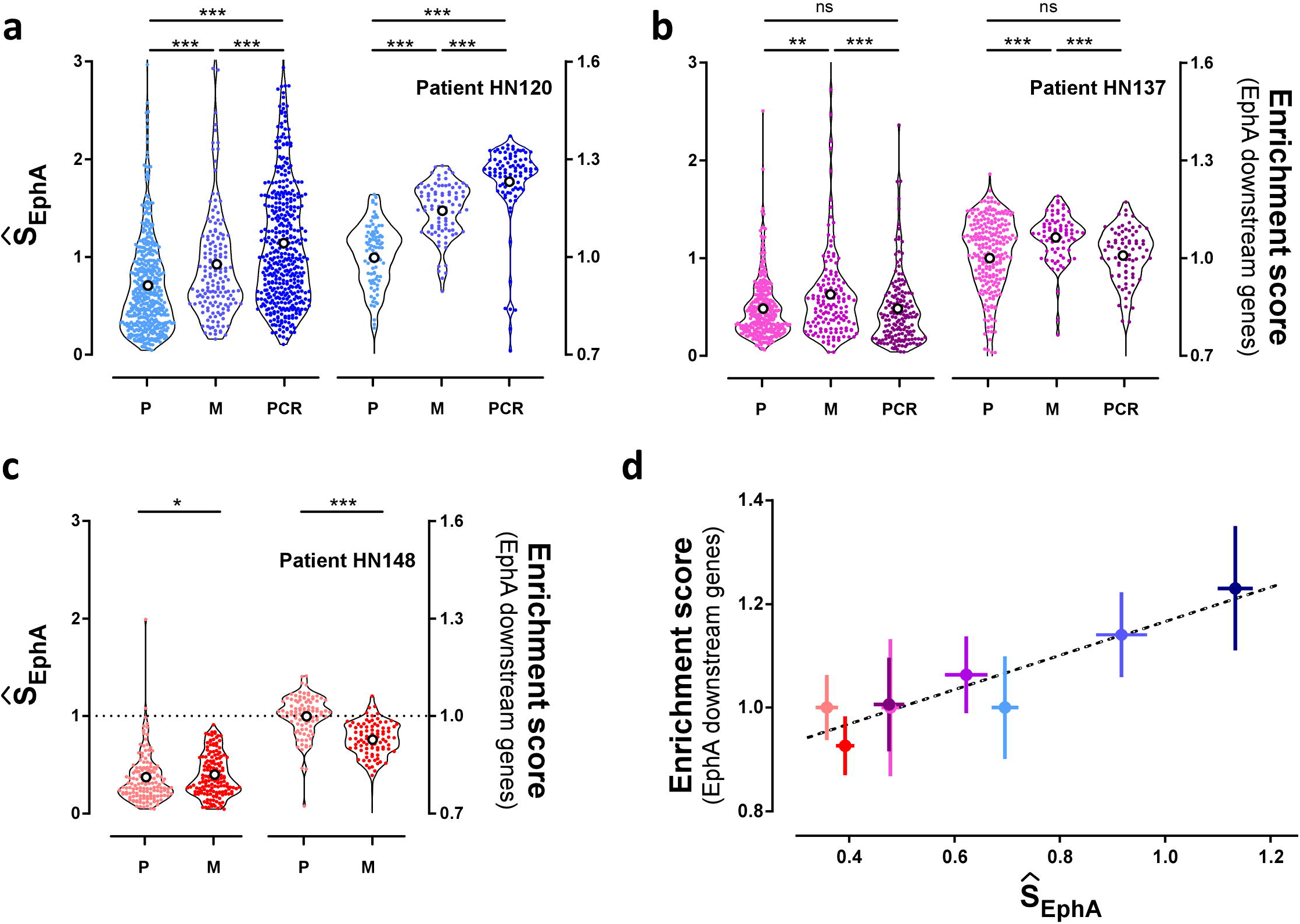
Comparison of single-cell phenotyping by EphA clustering for three different patients. **a)** Left axis: Distribution of 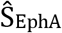 for patient-derived cell lines HN120 from patient 120. HN120P (P) was derived from a primary tumor, HN120M (M) from proximal lymph node, and HN120pcr (PCR) is an induced, cisplatin resistant cell line derived from HN120p. Right axis: Transcriptomic analysis (Enrichment score) obtained from the single-cell analysis of 71 genes positively correlated with EphA2 activation. Values are normalized by the average value of P. **b** and **c)** identical studies for patient 137 (HN137) and 148 (HN148). **d)** Correlative analysis of EphA clustering 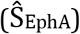 and EphA transcriptomic downstream signalling (Enrichment score). Strong correlation (Pearson coefficient = 0.93, p =0.001) between 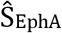 and transcriptomic scores demonstrate correspondence between EphA clustering and EphA downstream signaling.

Our results from Patient HN137 (Figure 7b) showed that HN137p and HN137pcr cells did not display any significant differences in 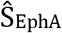 scores. However, scores in HN137m cells were slightly higher than that observed in HN137p, in line with the observed invasion potentials [22,24]. A similar trend was reflected in the transcriptomic scores. Patient HN148 (Figure 7c) displayed equal scores in HN148p and HN148m cells, although the transcriptional scores were slightly smaller in HN148m cells than in HN148p cells. Furthermore, 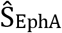 scores and transcription score displayed strong correlation (Pearson coefficient = 0.93, p =0.001), thereby indicating that our clustering score provides a reliable measure for EphA associated gene expression profile and phenotypes.

Overall, single-cell analysis of EphA clustering behavior and transcriptomic response in PDCs showed that our assay provided a reasonable assessment of the evolution of cell-states within these patient samples in a fast and cost-effective manner.

## Discussion

Intra-tumor heterogeneity poses a major challenge in therapeutic outcomes for cancer treatment in the clinic. Different clonal or sub-clonal cell populations can exhibit differential response to a given drug, and utilize distinct mechanisms to evolve therapy resistance, such as cell-state change like EMT or dormancy induced quiescence. Therefore, it is important to develop methodologies that can probe this heterogeneity to be able to identify novel genetic and therapeutic vulnerabilities. While existing single cell genomic/transcriptomic platforms have been shown to be extremely powerful in defining intra-tumor heterogeneity and the identification of new cell types and cell states, methodologies that can uncover functional heterogeneity at the level of single cells remain largely under-explored. In our study, we show that EphA cluster morphologies, which are associated with distinct migratory state of cells, constitute a single-cell-inherited trait of cell populations. We developed a common analysis scheme to quantitatively correlate cluster morphologies with the intrinsic migration potentials of cells. Additionally, results from this study extend to multiple cancer types our previous conclusions based on breast cancer cell lines [15] that cluster morphologies are indicative of the degree of transcriptional activation of genes involved in EphA signaling cascade. This signaling pathway largely involves cytoskeleton regulatory genes, which can explain the correlation between cluster morphologies and migration properties. By using PDCs from various cancer types, our study reinforces the growing body of evidence [13–16] that clustering properties of juxtacrine receptors can be used to define the “phenotypic state” of single cells. We conclude that this assay provides a technically simple, cost-effective and effective strategy to follow the gradual buildup of intracellular heterogeneity either upon cell division, (Figure 5) or across genomic drifts and drug selection, (Figure 7).

The rationale to develop parallelizable phenotypic assays stems from the observation that expression levels of genes related to a certain pathway do not always directly correlate with the behavioral response of cells. It is therefore important to develop coast-effective and simple behavioral assays that allow phenotypic probing of response to an activated pathway. Here we describe an assay based on EphA/ephrinA1 ligand signaling. Other membrane bounds RTK ligands could also be utilized in a similar fashion [25] provided that they prompt adhesion of the cells onto the bilayer. Such an approach is nonetheless restricted to ligands that foster cell adhesions and spreading on fluid bilayers. For other ligands, alternative approaches such as coupling several ligands together, could be devised [26].

Currently, most phenotypic assays rely on assessment of migration based on cell interactions with extracellular proteins. Compared to migration assays, our approach is fast and relevant at the level of single cell. It provides an easy and accurate assessment of functional intra-tumor heterogeneity in response to EphA activation, which is also indicative of the heterogeneity in migration potentials of single cells within a population. Future, parallelization of sample acquisition and development of automated acquisition and analytical platform would allow implementation of the current assay into a novel diagnostic tool to interrogate the heterogeneity and evolution of cancer states in clinical samples.

## Methods

### Preparation of Small Unilamellar Vesicles (SUV)

A clean, 5 ml round-bottom flask (PYREX) was rinsed with 100 % chloroform (Acros Organics) twice. A small amount of 100 % chloroform was left in the flask after rinsing. To generate SUVs using the sonication approach, 96 mol % 1,2-Dioleoyl-sn-glycero-3-phosphocholine (DOPC; Avanti Polar Lipids, cat. no. 850375) and 4 mol % 1,2-Dioleoyl-sn-glycero-3-[(N-(5-amino-1-carboxypentyl)imidodiacetic acid)succinyl] (nickel salt) (DOGS-NTA-Ni; Avanti Polar Lipids) were mixed in chloroform in the flask. A water bath apparatus from a rotary evaporator (EYELA, Japan) was warmed at 50º C to prevent the solvent from freezing during the evaporation process. The mixture of liposome and NTA-Ni was dried, first by using the rotary evaporator and then under a gentle stream of N_2_ for about 1 min, to prevent the exposure of lipid film to air. Dried lipids were resuspended in 1 ml of ultrapure (Milli-Q) water at room temperature as milky suspensions of multilamellar vesicles. To release physically adsorbed lipids from the walls of the flask, the flask containing the lipid solution was immersed in a warm water bath (60º C) for 10 sec. The lipid suspension was then transferred to a round-bottom polypropylene tube and sonicated for 20 min until the suspension became clear. After sonication, the clear lipid suspension was transferred to a 1.5 ml Eppendorf tube and centrifuged at maximum speed at 4º C, for at least 4 h. Finally, the supernatant containing small unilamellar vesicles was aliquoted and stored at 4º C.

### Preparation of devices made of cover glass and silicon chambers

Microscope cover glasses (22 × 22 mm) from High Precision were carefully cleaned to remove impurities by using a series of washing steps. Briefly, cover glasses were washed with dish-washing soap and rinsed with tap water. Thereafter, they were placed in a ceramic rack and sonicated in a sonicator bath (Elma) for 10 min, submerged in 100% acetone (Fisher Chemical). After rinsing with Milli-Q water, they were sonicated for 30 min, submerged in 1:1 isopropanol/Milli-Q water solution (isopropanol from Fisher Chemical). After thorough rinsing with deionized water, the cover glasses were incubated overnight submerged in a 1:1 sulfuric acid/Milli-Q water solution (sulfuric acid from Sigma) to remove all traces of organic solvent. Finally, they were rinsed with Milli-Q water, blow-dried using nitrogen gas, and placed in an 80º C dry oven for at least 2 h. Meanwhile, a PDMS silicon foil of 0.25 mm thickness (STERNE SAS, France) was cut into square chips of 20×20 mm sizes and further subdivided into 9 recording chambers (3 × 3 mm) using a cutting plotter (Graphtec Corporation, Japan). Importantly, one of the corners of the square was also cut to break the double symmetry and allow easy identification of wells (see Figure 1a and **Supplementary Figure 1a** and **b**). As the PDMS foil was polymerized in between two plastic films, the PDMS cuts were free from dust and contaminants. Finally, PDMS cuts were mounted on the glass coverslips and bounded using air plasma for at least 15 min at high power in a plasma cleaner (Harric Plasma). This step also provided a final cleaning of the surfaces and activation of the glass to facilitate adsorption of the lipids bilayer as described in the next paragraph.

### Preparation of ephrinA1 functionalized supported bilayer

The clean coverslip was placed on the lid of a 35mm petri dish with the PDMS gasket facing up. 4 µl of SUV TBS solution (1:1 ratio of SUV and 2× TBS solution, TBS form Sigma) was placed in each hole of the device. Upon contacting the activated surface of the plasma-treated cover glass, SUV spontaneously adsorb and formed a supported bi/multi-layer. From this point onwards, care must be taken to avoid direct contact of the supported bilayer with any air-liquid interface. After incubation for 5 min at room temperature, the device was submerged in a large beaker containing Milli-Q water and it was washed by vigorous sideways shaking to remove excess phospholipids from the bilayer. Thereafter, the device was placed in a 35 mm petri dish containing 1 ml of Milli-Q water. 1ml of 1 % BSA (Sigma) dissolved in 2× TBS solution was then added to the dish to block the unspecific binding of cells onto the glass or the PDMS gasket (final concentration of BSA about 0.5 %). The supported lipid membrane on the device was incubated in the blocking solution on a rotary shaker (gentle rotation) for 30 min at room temperature. After incubation, the device was washed 3 times with 1× TBS solution, and was then left submerged in 2 ml of 1× TBS solution. Meanwhile, ephrinA1 protein was purified and conjugated with Alexa 568 with NHS ester according to the protocol we developed in [15]. It was dissolved in 1× TBS solution at a dilution ratio of 1:2000. 100 µl of protein solution was then added to the device in the petri dish. The device was then incubated with protein solution for 1 h on a rotary shaker (gentle rotation) at room temperature, in the dark. Conjugation of protein to the supported bilayer occurs during this step. After incubation, excess protein solution on the membrane was washed using imaging buffer (150 mM NaCl, 30 mM KCL, 2 mM Ca2Cl, 2 mM Mg2Cl, 10 mM Hepes, 10mM D-Glucose, pH 7.4). The device was then placed in a petri dish and submerged in 2 ml of imaging buffer. Devices were freshly prepared and long-term storage was avoided. Devices were kept covered in the dark at room temperature while waiting for the cells to be seeded. The devices were acclimated inside a 37º C incubator for at least 1 h before cell seeding.

### Cell culture, dissociation, seeding and fixation

All the cells used in this study were tested for mycoplasma using Mycoprobe Mycoplasma Detection kit from R&D system. Cells were passaged no more than 10 times from the initial stocks. The lung, ovarian, gastric and breast cancer cell lines were obtained commercially. The isolation and derivation of the PDCs are extensively described in [22]. Briefly, tumors were minced and dissociated enzymatically using 4mgmL−1 collagenase type IV (Thermo Fisher, cat. no. 17104019) in DMEM/F12, at 37 °C for 2 h. The cell suspensions were strained through 70 μm cell strainers (Falcon, cat. no. 352350), prior to pelleting and resuspension in RPMI (Thermo Fisher, cat. No. 61870036), supplemented with 10% foetal bovine serum (Biowest, cat. no. S181B) and 1% penicillin-streptomycin (Thermo Fisher, cat. no. 15140122). The identities of the cell lines were checked comparing the STR profile (Indexx BioResearch) of each cell line to its original tumor. A standard cell culture incubator at 37º C with 5 % CO_2_ was used to expand all cells. The cells were sub-cultured at 80 % confluence in T25 flasks (Thermo Fisher Scientific) and reseeded into fresh flasks at approximately 30-50 % confluence. Prior to the experiments, cells were grown at 50 – 70 % confluence and dissociated with an enzyme-free dissociation buffer (Gibco) for about 15 to 30 min. In the meantime, the device was kept submerged in imaging media in a 35 mm petri dish and was acclimated inside an incubator at 37º C. After removing the imaging buffer from the petri dish, the chambers of the device were left filled with imaging media (approx. 2 µl). Care was taken in this very critical step to avoid exposing the lipid bilayer to air-liquid interface at all times. Immediately after removal of the media, 5µl of cell suspension was directly deposited on top of each chamber of the device, forming small drops with high contact angle due to the hydrophobicity of the silicon-based device. Spacing between the chambers kept cells contained within their respective chambers. Seeding in alternate chambers ensured the absence of overspill of cells from adjacent chambers. The three cell types were labeled with different fluorescent markers (stably transfected GFP and RFP cell lines and one with Hoechst live nuclear stain). Due to the low volume and height of the chambers, seeded cells nearly immediately contacted the bilayer. About 200µl of sterile Milli-Q water was pipetted along the walls of the petri dishes to provide extra humidity and prevent evaporation of water from the chambers. Petri dishes were placed in a secondary container (typically a 140 mm petri dish with humidifying pads soaked in sterile Milli-Q water) previously acclimated at 37º C. The secondary container was then placed inside a 37º C incubator for 30 min to allow EphA clustering to occur at nearly physiological conditions. After 30 min, clustering was stopped by PFA fixation: devices inside the 35 mm petri dish were flooded with 2 ml of 4 % PFA solution (Thermoscientific). Cells were fixed at 37º C for 15 min. Thereafter, devices were washed with 1× PBS solution for 5 times and incubated in 10 mM NH_4_Cl solution for 10 min at room temperature. Devices were then washed with 1× PBS solution for 3 times and mounted on glass slides using fluorescent mounting medium (Dako) and the coverslips were sealed using nail polish. Short-term storage of the devices was done at 4º C overnight to a maximum of 2 days before imaging.

### Microscopy and image analysis

Fixed samples were imaged using an IX81 microscope (Olympus) equipped with an 100X oil-immersion objective (UAPON 100XTIRF, NA 1.49, Olympus), X-Cite light source (Excelitas Technologies), a CoolSNAP EZ CCD camera (Photonics) and an appropriate filter set to detect Alexa 568 conjugated to ephrinA1. Pairs of bright-field and fluorescence images were acquired for each field of view. The focus of the objective was adjusted to visualize the supported bilayer. Images were analyzed using a custom MATLAB code (can be provided upon request). The code performed bunch analysis of images found in the input folder. The analysis was divided into two separate steps: First, cell segmentation was performed by manually outlining cell borders on the composite image generated from the bright-field and corresponding fluorescence images. This process defines the regions of interests for subsequent analysis and generates a third masking image where the ROI are coded with ascending natural numbers. In the second step, image intensity distribution from the ROIs was analyzed and the image entropy for each cell quantified according to *S*_EphA,_ = ∫ *ρ* ln *ρ*, where 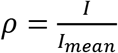.

### Western Blotting

Protein lysates (30 µg) from individual cell lines were firstly dissolved in SDS-PAGE loading buffer and loaded onto 10% SDS-PAGE gel. Using Bio-Rad Mini-PROTEAN electrophoresis system, protein samples were run at 110V, until samples ran completely through the stacking gel and into the resolving gel. The voltage was then increased to 150V, until dye front ran off the bottom of the gel. After protein separation by SDS-PAGE, proteins were transferred from gel to a PVDF membrane (Merck, cat. no. IPSN07852). Transfer was carried out for 45 minutes at 110V in transfer buffer (cold room). Following this, the PVDF membrane was blocked with 5% skim milk powder in TBS-Tween20 (0.1% v/v) at room temperature for approximately 1 h. Anti-EphA2 (Cell Signaling, cat no. 6997) and anti-beta actin (Pierce, cat. no. MA515739) were used as primary antibodies and diluted in blocking solution. The membrane was first probed with primary antibodies at 4ºC overnight. Anti-rabbit HRP (Invitrogen, cat. no. G21234) and anti-mouse HRP (Invitrogen, cat. no. G21040) were used as secondary antibodies and diluted in blocking solution. The membrane was then probed with secondary antibodies at room temperature for 1 h. The membrane was washed 3 times for 10 min in TBS-Tween20 (0.1% v/v) on an orbital shaker after blocking with milk and after each probing step. Finally, the membrane was treated with chemiluminescent substrate (ThermoScientific, cat. no. 34080) and protein bands were detected by Bio-Rad Chemidoc imaging system.

### Wound-model assay

Classical wound-model assays have been performed [27,28]. Briefly, a rectangular cut of PDMS was placed in the middle of a 6-well plate and let to adhere for 2 h at 37º C before cell seeding was performed. Cancer cells were seeded around the PDMS cut and allowed to grow to full confluence inside a cell culture incubator for about one day. After removal of the PDMS cut, the 6-well plate was placed inside a Biostation CT (NIKON) and time lapses of cell migration by phase contrast were acquired for 12 h (time interval between image acquisitions was 15 min). Cell movement within the first 50 µm from the wound edge was analyzed using Particle Image Velocimetry in MATLAB.

### Single-cell transcriptomic analysis

We generated single-cell gene expression matrix for 682 cells from patient derived cell lines (PDCs) from head and neck (HN) carcinomas from primary site (P), secondary/metastatic tumor (M) and primary with acquired cisplatin resistances (PCR) from three patients (patients 120, 137 and 148). Thus, PDC nomenclature is HN120p, HN120m, HN120pcr, HN137p, HN137m, HN137pcr, HN148p and HN148m [22]. The Single cell RNA-seq datasets is available in the Gene Expression Omnibus repository, at the link: https://www.ncbi.nlm.nih.gov/geo/query/acc.cgi?acc=GSE117872. Expression levels were quantified as E=log_2_(TPM+1). We selected only the subset of genes putatively involved in the EphA2 signaling cascade [15]. Potential markers with average expression levels lower than 1 across all the cells were excluded. We defined single-cell genomic score as the linear average of the expression levels of the remaining genes. Transcriptomic scores of each patient were normalized by average value of primary samples to account for individual genetic background. Five cells with low EphA2 score (<3) were discarded as outliers.

## Supporting information

Suppl.Mov1

Suppl.Mov2

## Acknowledgements

V. Viasnoff and J.T Groves acknowledge the support of the collaborative NRF CRP11-2012-02 grant, and MBI seed funds. A. Ravasio and C. Bertocchi acknowledge financial support from seed founds of the Pontificia Universidad Católica de Chile. R. Dasgupta acknowledges funding from partly the National Medical Research Council (Singapore) (NMRC/CIRG/1434/2015). Proof reading was kindly provided by Andrew Wong and Sruthi Jagannatha. We would like to extend our gratitude to all patients and families involved in this study.

## Authors contribution

AR, RYJH, RDG, JTG and VV conceived and designed the study. AR, MZM, SC, AA, CB, BA, ACG, and ZC collected and analyzed the data. SC, EYC, VYC, TZT, RYJH and RDG designed and performed transcriptomics analysis. AR, CB, HTO, JTG and VV designed and performed image analysis. AR and AnSa designed and performed machine-learning method. SC, AnSk, OGI and RDG produced patient derived and drug resistant cell lines. AR, MZM, CB, RDG, and VV wrote the paper. All authors discussed the results and contributed to the final manuscript.

