## Supplementary material for "Single-cell analysis of EphA clustering phenotypes to probe cancer cell heterogeneity": Suppl.Info

[1] Mechanobiology Institute, National University of Singapore, Singapore, Singapore, [2] Pontificia Universidad Católica de Chile, Santiago, Chile, [3] Genome Institute of Singapore, A-STAR, Singapore, Singapore, [4] Cancer Science Institute of Singapore, National University of Singapore, Singapore, [5] National Cancer Centre, Singapore, [6] University of California, Berkeley, CA, USA, [7] Centre National de la Recherche Scientifique, Singapore, Singapore.

**Supplementary Material:**

**Image feature extraction and unbiased machine learning method**

Image features of fluorescence images were extracted using custom python code (open code at: https://github.com/aneeshsathe/Ephrin_cluster_analysis) Briefly, cell boundaries were manually identified from brightfield images (see image analysis in main text) and used to generate a mask identifying the region of interest (ROI). ROI were used on fluorescent images for feature extraction. We used the scikit-image pacakge [1] to extract commonly used features. This list was further populated by implementing simple modifications/normalization of the features provided in the scikit-image package. A total of 75 image features accounting for texture, morphology, and intensity properties of the EphA clustering were considered (**Supplementary Table 1**). Custom code was written to extract custom and gray level run length matrix (GLRLM) [2] features.

To visualize the distinct groups in the extracted features we used t-Distributed Stochastic Neighbor Embedding (t-SNE) (**Supplementary Figure 2a**). The t-SNE plot was generated using the embedding generated by linear discriminant analysis(LDA) with the scikit-learn [3] python package.

The machine learning method, random forests [4] was used to train a classifier for the different cell types. Using the 75 features, the model predicted the correct cell type with a ~70% accuracy. This model was used to then extract the relative important of each feature to identify different cell types(**Supplementary Figure 2b**). The confusion matrix for the random forests model is in **Supplementary Table 2**.

**Statistical relevance of the single colonies.**

The number of cells that can be analyzed from clonal after 10 days is intrinsically limited by the proliferation rate of the cells. For HN137p 10 days corresponded to an average of 10 divisions. This estimation was based on measuring the colony area as well as the average cell area. This crude estimate did not lead to any significant variation of growth rate between the different colonies. After ten 10 the colonies have an average of 1200 cells. Splitting the colonies to run a fraction on the assay and keep growing the rest for another 10 days allowed us to analyze only **100** cells per colonies. We had to quantify the likelihood that the measured distributions contains a clear signature of the original single cells from which we grew the colony. To this end we estimated the likelihood that the measured distribution for each colony stems from the random picking of 100 cells from the whole population distribution measured for HN137p cells.

First we created two "random" distributions $P_1$ and $P_2$ of 100 cells by randomly picking 100 values from the whole population distribution. We calculated the minimal Euclidian distance d between the two distributions by sorting in ascendant order the values of each distribution ($P_1^s, P_2^s$) and calculating the norm of the resulting vector difference.

$$d = |P_1^s - P_2^s| = \sqrt{\sum_{i=1}^{100}[(P_1^s[i])^2 - (P_2^s[i])^2]}$$

For a given $P_1$ distribution the process was repeated $10^4$ times and the average distance of $P_1$ to any other 100 sample distribution was computed.

$$\langle d \rangle = \frac{1}{10^4} \sum_{P_2^S} d$$

Repeating the calculation $10^5$ times, we established the distribution $\Pi$ ($\langle d \rangle$ ) of the probability for a "random 100 sample" distribution to have an average distance $\langle d \rangle_P$ to any other "random 100 sample" distribution.

We then considered the distributions of $S_{EphA}$ measured for the clonal colonies $P_{clonal}$. We calculated the distance $\langle d \rangle_{P_{clonal}}$ and we estimated the likelihood of randomness of the measured distribution by the value $\Pi$ ( $\langle d \rangle_{P_{clonal}}$ ). For all the measured clonal distributions the likelihood of randomness is below 1%. The evaluation of statistical significance is illustrated in Supplementary figure 5.

**Supplementary Figures**

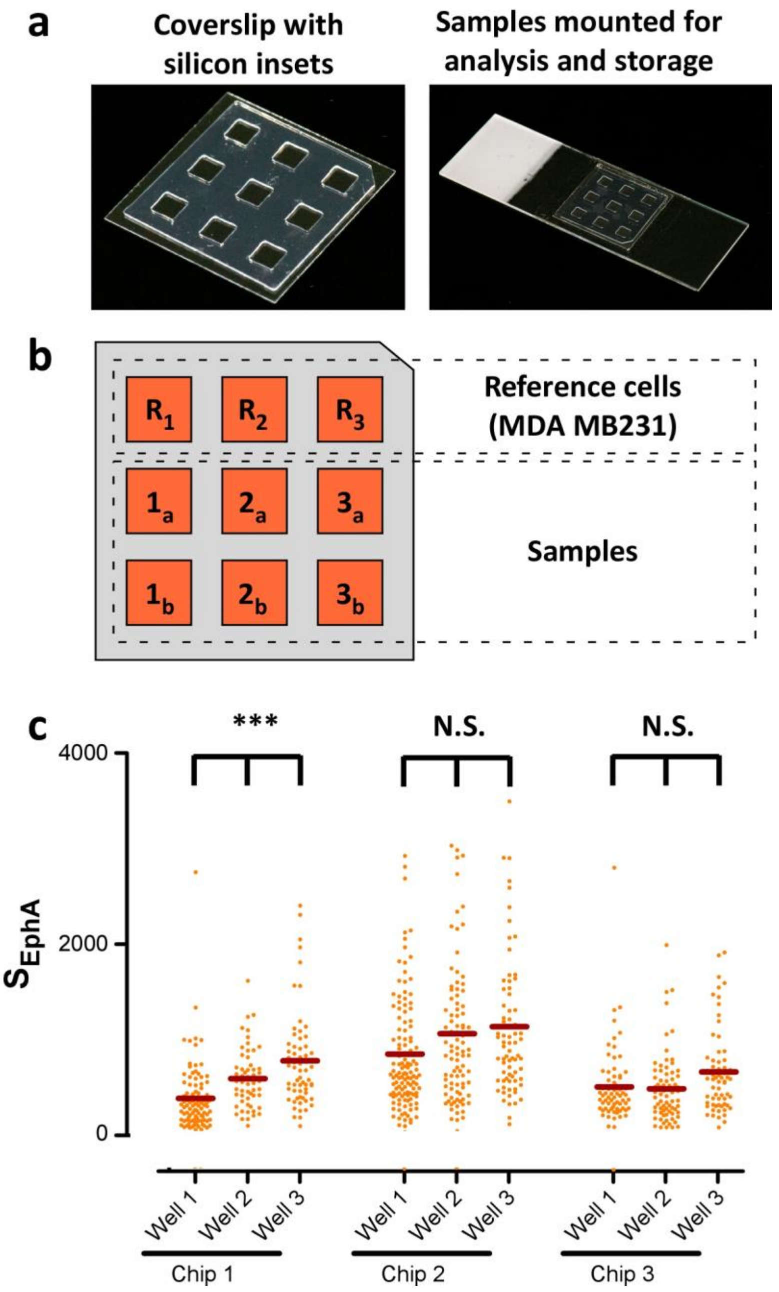


**Supplementary Figure 1 - Implementation of the assay a) Left**, imaging device: silicon gasket with 9 chambers of 5x5 mm mounted on glass coverslip. Dimensions of the wells have been optimized to obtain a fluid bilayer (fluidity tested by FRAP – i.e. Fluorescence Recovery After Photobleaching experiments) able to contain about 50 to 100 cells per well. **Right**, sample mounted on a glass slide for analysis and storage After exposure to the biofunctionalized lipid bilayer, cells were fixed with paraformaldehyde, submerged in mounting media and mounted on a glass slide using transparent nail polish. Microphotographs of the samples have been acquired within 2 days after mounting. Samples could be stored up to 2 weeks at 4°C in a dark box with wet patches. **b)** Arrangement of samples and references within the device. For each device, 3 wells were used to systematically quantify clustering of reference cell lines MDA MB231 in triplicate (**R**$_{1\text{-}3}$). This cell line served as internal control for the quality of the device and as reference to normalize absolute $S_{EphA}$ values. Remaining wells were used to test two samples (marked as **a** and **b** subscripts) in triplicates (well **1** to **3**). Image acquisition of the samples was performed following a vertical raster mode (from R1 to 1a to 1b). **c)** Intra and inter device variability of the measured distribution of $S_{EphA}$. ANOVA analysis of the triplicates with threshold of 0.05 (Kruskal-Wallis non-parametric test and Dunn's post-test multiple comparison) was used to identify devices with defective bilayers. Chips variability above the threshold indicates for defective supported bilayers (e.g. **chip 1** of this series was discarded). Only chips presenting a non-significant intra-chip variability for MDA MB231 are considered (**chip 2** and **3**). They constitute >80% of the chips. Thereafter, $S_{EphA}$ values were pulled together and their average was used to normalize the $S_{EphA}$ values of the samples measured in the same chip. Normalized values are noted as $\hat{S}_{EphA}$ (see fig. **3b** and **c** in the main text).

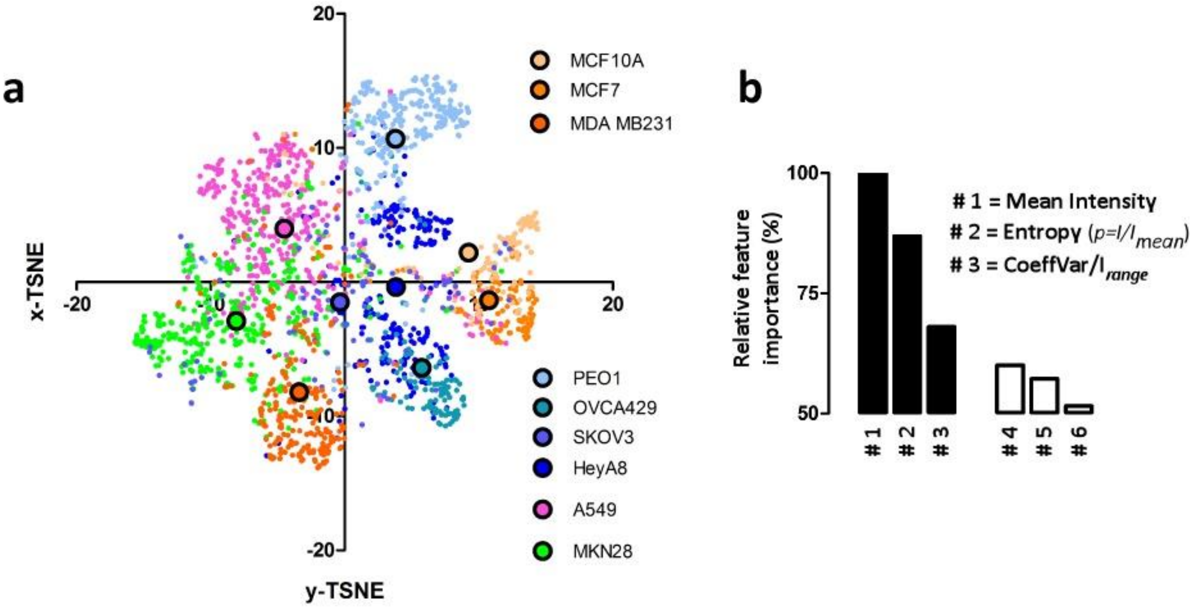


**Supplementary Figure 2 - Unsupervised clustering of 12 cell types based on EphA cluster morphologies (72 features tested) a)** Feature extraction and machine-learning was used to identify discriminatory features from the EphA. Data representation using TSNE demonstrates separation of cell types based on 75 image features derived from morphology, texture and intensity. **b**) Machine learning (random forests) was used to build a cell type classifier based on the image features (accuracy of cell type identification >70%). Top 3 image features contributing the most to identify and differentiate cell types are **mean intensity**, modified **image entropy** and **coefficient of variation** normalized by the intensity range. **Mean intensity** was discarded because it reports extensive properties of the system (amount of ligand engaged, fluorophore quantum yield, illumination, gain of the camera. etc). The second-best classifier is the modified **image entropy**. This image feature is extensive and reports the intrinsic complexity of the image intensities in a localization independent manner (**Supplementary Figure 4**).

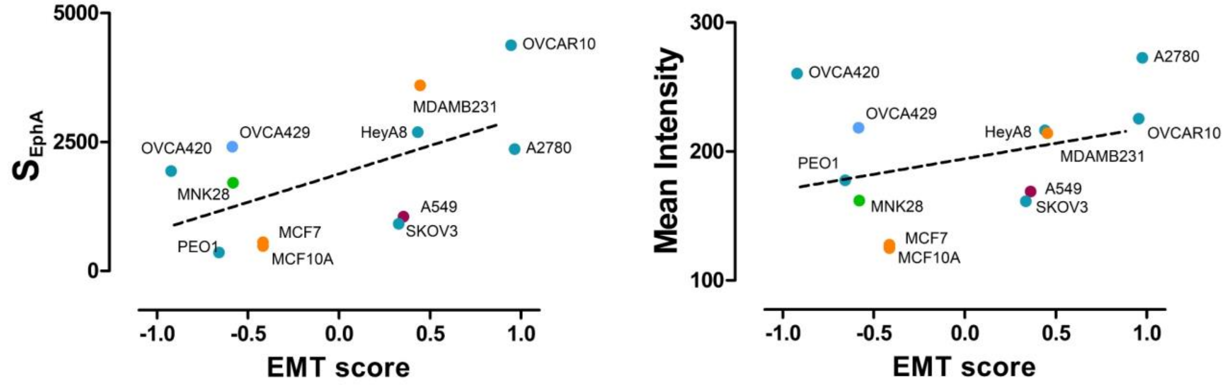


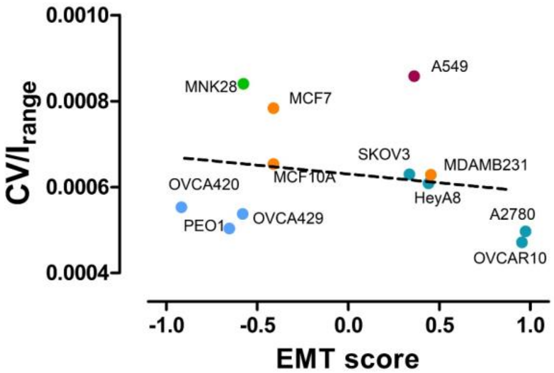


| | Pearson coeff. (r) | P value |
|---|---|---|
| $S_{EphA}$ | 0.56 | 0.05 |
| Mean Int. | 0.33 | 0.29 |
| $CV/I_{range}$ | -0.20 | 0.52 |

**Supplementary Figure 3 - Correlation between EMT scores and 3 candidates to score EphA cluster morphologies.** Correlative graphs and respective Pearson's coefficient table for Entropy, Mean Intensity and the normalized coefficient of variation. The top three discriminators from Random Forest analysis (**Supplementary Figure 2b**) are here plotted against the Epithelial to Mesenchymal Transition (EMT) score. EMT score reports the degree of genetic transformation of each cell line from epithelial to mesenchymal genotypes. Image entropy ($S_{EphA}$) best correlated with EMT.

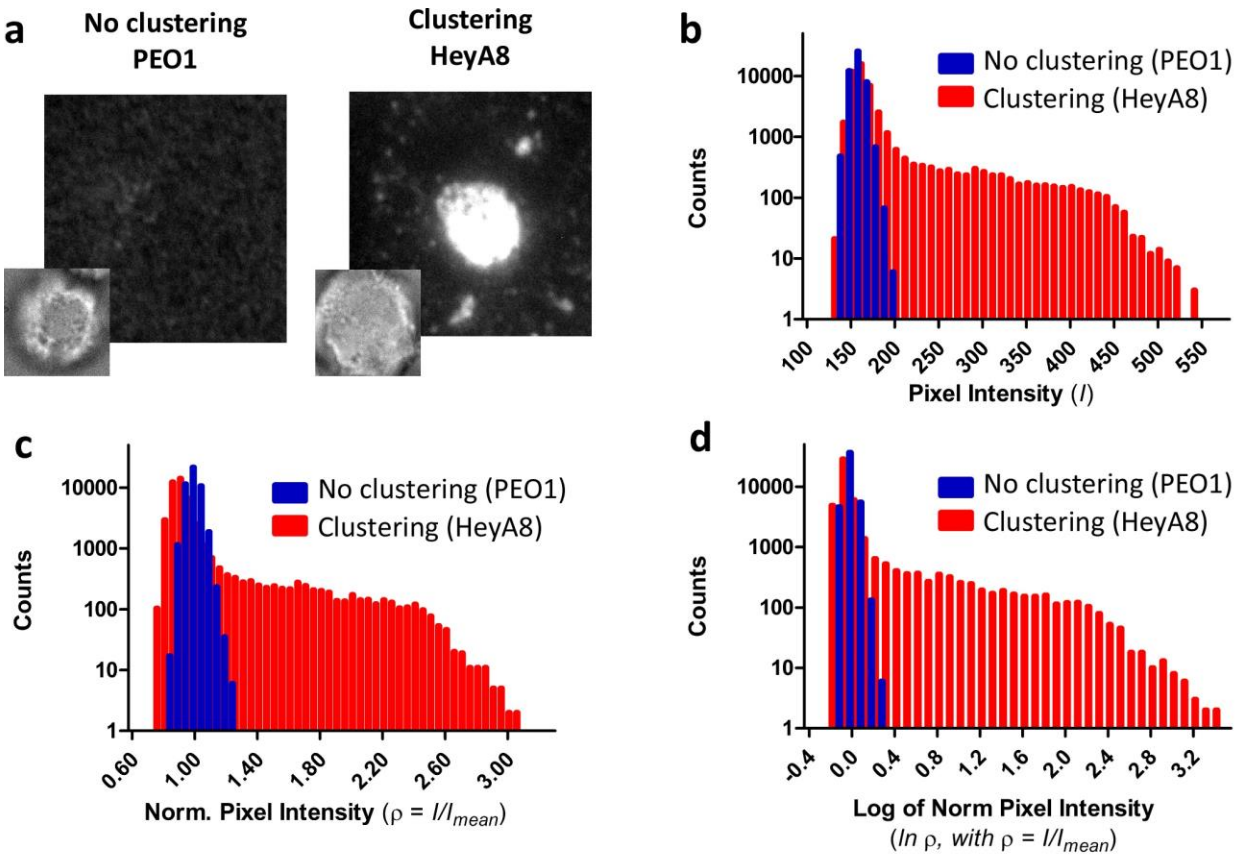


**Supplementary Figure 4 – Statistical distribution of fluorescence intensity values and modified image entropy. a**) Exemplary images from a cell without clusters (PEO1) and another displaying clustering patterns (HeyA8). Large micrographs show fluorescence images of ephrin A1. Small insets display the corresponding bright fields. **b**) Frequency distribution of fluorescence intensity values from images in panel **a**. When cells do not form clusters the distribution of the intensity values approximate a random Gaussian distribution. In contrast, an asymmetric nonrandom distribution appears in the form of a right tail in the presence of clustering. **c**) Values in **b** normalized by their respective mean values define a modified version of the probability states ($\rho = \frac{I}{I_{mean}}$). **d**) Logarithm of values in **c** ($\ln \rho$). Thus, the function $S_{\mathrm{EphA,}} = \int \rho \ln \rho$, where $\rho = \frac{I}{I_{mean}}$, tends to zero in the absence of clustering (positive and negative values for $\ln \rho$ (**d**) are nearly the same) and it becomes grater as clustering of the protein progress (shift toward the right of intensity distribution).

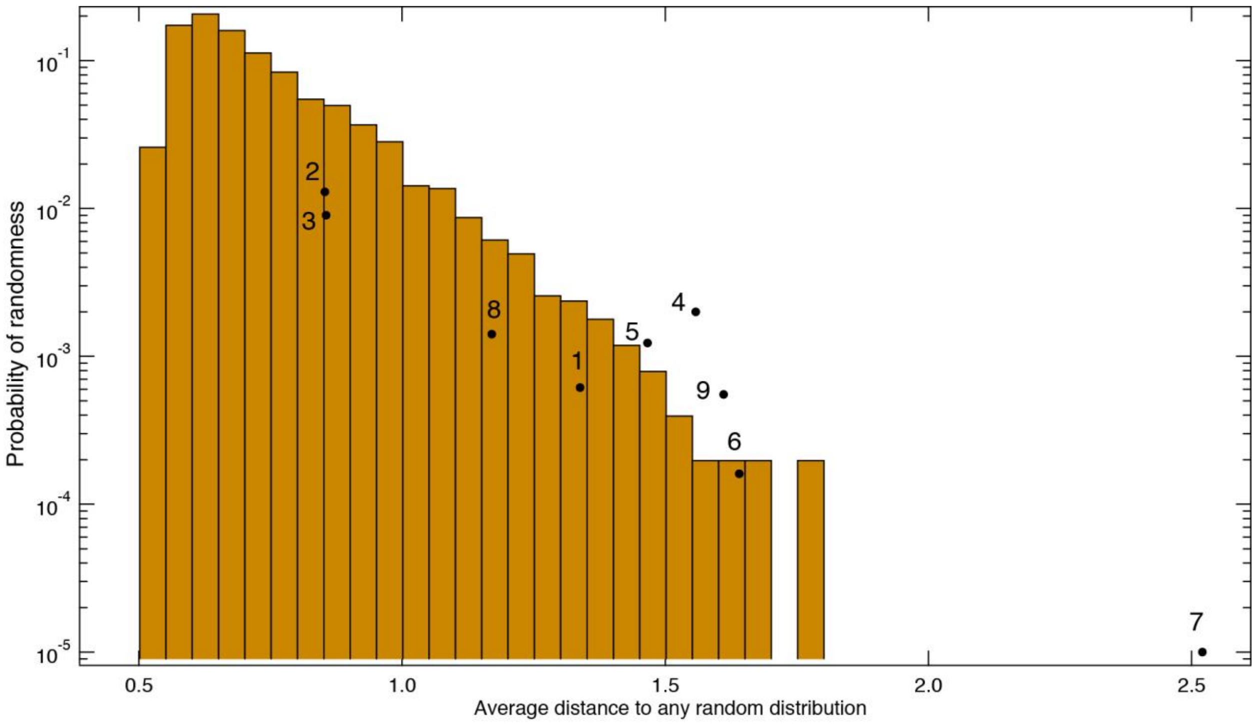


**Supplementary Figure 5 – Assessment of the probability of randomness of the distribution of scores obtained for single colonies.** Sub-distributions containing 100 cell scores were generated by random picking from the 1200 values of the distribution for the whole cell population. We repeated the operation to establish the probability of Euclidian distance between two of these sub-distributions. This probability distribution is represented on a logarithmic scale in maroon. WE then assessed the average distance (dots) of the score distributions obtained for clonal colonies with any random sub-distribution. Each dot is labeled by the colony number. We found that all our clonal colonies display a probability of randomness below 1% and 6 colonies are below 0.5% of randomness probability. It demonstrates that the distribution of scores obtain for each company does not reflect the scattering of scores with the whole population but is directly related to the score of the original cells from which the clonal colony is derived.

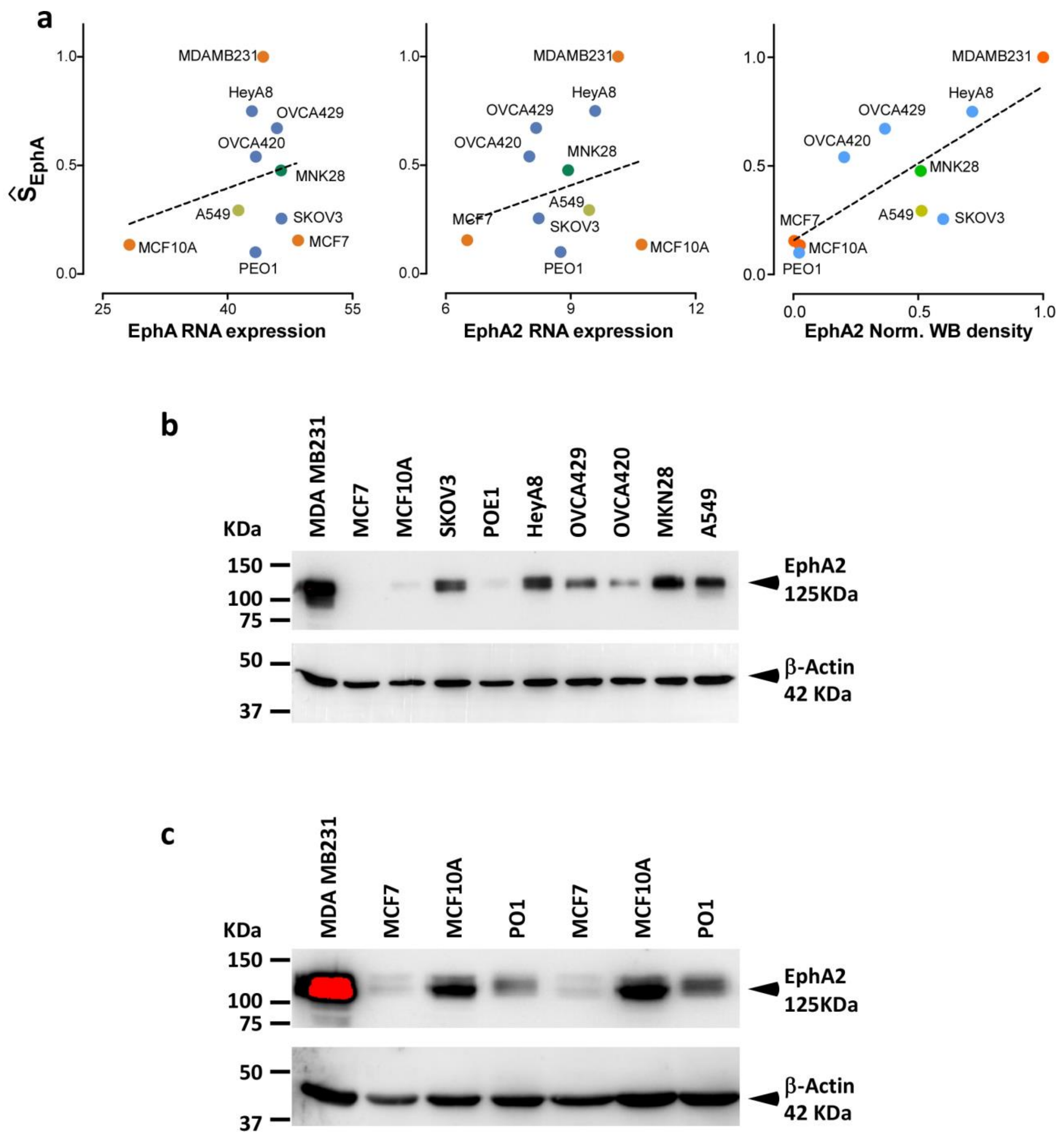


**Supplementary Figure 6 – Western Blot and quantitative measurement of the EphA2 expression level. a**) Plot correlating the average $\hat{S}_{EphA}$ value for each cancer population with the EphA mRNA expression (left, Pearson coefficient = 0.26, p = 0.47), EphA2 mRNA (center, Pearson coefficient = 0.27, p = 0.46) and EphA2 protein (right, Pearson coefficient = 0.97, p =0.007). **b**) Exemplary Western blot for 10 different cell lines (n=3). **c**) More sensitive WB (higher concentration, larger exposure time) to analyze and quantitate the cells with very low expression levels (e.g. MCF7).

**Legend of Supplementary Movie 1 and 2**

Time-lapse of EphA cluster formations in PEO1 cell (**Supplementary Movie 1**) and HeyA8 cell (**Supplementary Movie 2**). **Left**, fluorescence images of Alexa568-ephrin A1. Ligand has free later diffusibility on the supported bilayer. Thus, dynamics of clustering are to be attributed to active transport of the EphA receptor by the cell cytoskeleton. **Middle**, Reflection Interference Contrast Microscopy (RICM) images. Dark regions (destructive interference) denote the location where cell membrane is intimate contact with the supported bilayer. Importantly, these regions correspond to the highest fluorescence on the left images. **Right**, phase contrast images of the cells. Movies length = 30 min. Frame rate = 30 seconds/frame.

**Supplementary Tables**

**Supplementary Table 1 -** Table of image features

| | | |
|---|---|---|
| **1** | **Stdev1** | **std = sqrt(mean(abs(x - x.mean())**2))** |
| **2** | StdevByIntRange2 | stdev_byIntRange = stdev_int/(np.max(int_vec)-np.min(int_vec)) |
| **3** | CoeffVar3 | coeffvar_int = stdev_int/mean_int |
| **4** | CoeffVarByIntRange4 | coeffvar_byIntRange = coeffvar_int/(np.max(int_vec)-np.min(int_vec)) |
| **5** | StdevPerum5 | stdev_int_per_area = stdev_int/area_um |
| **6** | CoeffVarPerum6 | coeffvar_int_per_area = coeffvar_int/area_um |
| **7** | Entropy7 | entropy_orig = -scipy.stats.entropy(int_vec) |
| **8** | EntropyPerPix8 | entropy_per_pixel = entropy_orig/use_area |
| **9** | EntropyNew9 | p1 = int_vec/np.float(mean_int)<br>p = p1[p1 > 0]<br>entropy_new = sum(p*np.log(p)) |
| **10** | EntropyNewPerum10 | entropy_new_perum = entropy_new/area_um |
| **11** | EntropyNewBySumInt11 | entropy_new_by_sumInt |
| **12** | EntropyHist12 | pro_bin_prob, bin_cen = histogram(region_prop.intensity_image[region_prop.image])<br>bin_prob = np.float16(pro_bin_prob) / sum(pro_bin_prob)<br>bin_prob = bin_prob[bin_prob > 0] |

| | | |
|---|---|---|
|  |  | `entropy_hist = sum(bin_prob * np.log2(bin_prob)) # entropy` |
| **13** | EntropyHistPerum13 | `entropy_hist_perum = entropy_hist/area_um` |
| **14** | EntropyHistBySumInt14 | `entropy_hist_by_sumInt = entropy_hist/sum_int` |
| **15** | UniformityHist15 | `uniformity_hist = sum(np.square(bin_prob))` |
| **16** | Kurtosis16 | `out_kurtosis = scipy.stats.kurtosis(int_vec)` |
| **17** | Skewness17 | `out_skew = scipy.stats.skew(int_vec)` |
| **18** | Smoothness18 | `out_smoothness = 1.0 - (1.0 / (1.0 + np.square(np.std(int_vec))))` |
| **19** | SumInt19 | `sum_int = sum(int_vec)` |
| **20** | MeanInt20 | `mean_int = sum_int/use_area` |
| **21** | MinInt21 | `min_int = min(int_vec)` |
| **22** | MaxInt22 | `max_int = max(int_vec)` |
| **23** | Area23 | `region_prop.area, region_prop.eccentricity, region_prop.euler_number, region_prop.equivalent_diameter, region_prop.extent, region_prop.major_axis_length, region_prop.minor_axis_length, region_prop.orientation, region_prop.perimeter, region_prop.solidity` |
| **24** | Eccentricity24 | |
| **25** | Euler_Num25 | |
| **26** | Equi_Diam26 | |

| | | |
|---|---|---|
| **27** | Extent27 | |
| **28** | Maj_ax_len28 | |
| **29** | Min_ax_len29 | |
| **30** | Orientation30 | |
| **31** | Perimeter31 | |
| **32** | Solidity32 | |
| **33** | Roundness33 | `out_roundness = mahotas.features.roundness(region_prop.image)` |
| **34** | MorphArea34 | |
| **35** | MorphPerimeter35 | |
| **36** | MorphCompactness36 | |
| **37** | MorphAspectRatio37 | |
| **38** | MorphSolidity38 | |
| **39** | MorphRoundness39 | |
| **40** | MorphEccentricity40 | |
| **41** | MorphCenterOfMassShift41 | |
| **42** | IntSumInt42 | |
| **43** | IntMeanInt43 | |
| **44** | IntMinInt44 | |
| **45** | IntMaxInt45 | |
| **46** | IntRangeInt46 | |
| **47** | IntMedianInt47 | |
| **48** | IntPercentile1048 | |
| **49** | IntPercentile9049 | |
| **50** | IntIQtR50 | |
| **51** | IntMeanAbsDev51 | |
| **52** | IntRobustMeanAbsDev52 | |
| **53** | IntMedianAbsDev53 | |

| | |
|---|---|
| **54** | IntStdev54 |
| **55** | IntCoeffVar55 |
| **56** | IntQuartCoeffDispersion56 |
| **57** | IntEnergyInt57 |
| **58** | IntRMSInt58 |
| **59** | IntEntropyInt59 |
| **60** | GLRLMShortRunEmphasis60 |
| **61** | GLRLMLongRunEmphasis61 |
| **62** | GLRLMLowGreyLevelEmphasis62 |
| **63** | GLRLMHighGreyLevelEmphasis63 |
| **64** | GLRLMShortRunLowGreyLevelEmphasis64 |
| **65** | GLRLMShortRunHighGreyLevelEmphasis65 |
| **66** | GLRLMLongRunLowGreyLevelEmphasis66 |
| **67** | GLRLMLongRunHighGreyLevelEmphasis67 |
| **68** | GLRLMGrayLevelNonUniforminty68 |
| **69** | GLRLMGrayLevelNonUniformintyNormalised69 |
| **70** | GLRLMRunLengthNonUniforminty70 |
| **71** | GLRLMRunLengthNonUniformintyNormalised71 |
| **72** | GLRLMRunPercentage72 |
| **73** | GLRLMGrayLevelVariance73 |
| **74** | GLRLMRunLengthVariance74 |
| **75** | GLRLMRunEntropy75 |

**Supplementary Table 2 -** Confusion matrix for the random forests model

| Predicted | A549 | HeyA8 | MCF10A | MCF7 | MDAMB231 | MNK28 | OVCAwt | PEO1 | SKOV3 |
|---|---|---|---|---|---|---|---|---|---|
| **Actual** | | | | | | | | | |
| **A549** | 259 | 5 | 7 | 5 | 2 | 32 | 1 | 11 | 1 |
| **HeyA8** | 16 | 113 | 1 | 0 | 11 | 25 | 15 | 8 | 2 |
| **MCF10A** | 14 | 0 | 74 | 14 | 0 | 0 | 0 | 2 | 4 |
| **MCF7** | 10 | 0 | 17 | 75 | 0 | 0 | 0 | 0 | 1 |
| **MDAMB231** | 16 | 12 | 0 | 0 | 126 | 24 | 4 | 2 | 2 |
| **MNK28** | 41 | 6 | 2 | 1 | 11 | 222 | 2 | 10 | 4 |
| **OVCAwt** | 2 | 25 | 3 | 0 | 4 | 2 | 61 | 3 | 1 |
| **PEO1** | 10 | 6 | 1 | 0 | 2 | 7 | 3 | 209 | 0 |
| **SKOV3** | 15 | 2 | 7 | 3 | 5 | 19 | 6 | 4 | 18 |

**Supplementary Table 3 – Statistical analysis of figure 4.** Tables reporting statistical analysis for $\hat{S}_{EphA}$ distributions across cell types. Kruskal–Wallis test with Dunn`s multiple comparison test to evaluate differences between mean values in **Figure 4a** and Kolmogorov-Smirnov test to evaluate differences in frequency distributions between different cell lines in **Figure 4b**. P < 0.05, one star; P < 0.01, two stars; P < 0.001, three stars; P < 0.0001, four stars.

| Figure 4a | MCF7 | MDAMB231 | PEO1 | SKOV3 | OVCA420 | OVCA429 | A2780 | HEYA8 | OVCAR10 | A549 | MKN28 |
|---|---|---|---|---|---|---|---|---|---|---|---|
| MCF10A | ns | **** | ns | ns | **** | **** | **** | **** | **** | **** | **** |
| MCF7 | | **** | ** | ns | **** | **** | **** | **** | **** | * | **** |
| MDAMB231 | | | **** | **** | **** | **** | ns | **** | ns | **** | **** |

| | | | | | | | | | | | |
|---|---|---|---|---|---|---|---|---|---|---|---|
| PEO1 | | | | **** | **** | **** | **** | **** | **** | **** | **** |
| SKOV3 | | | | | **** | **** | **** | **** | **** | ns | **** |
| OVCA420 | | | | | | ns | **** | ns | **** | **** | ns |
| OVCA429 | | | | | | | ns | ns | **** | **** | ns |
| A2780 | | | | | | | | ns | * | **** | **** |
| HEYA8 | | | | | | | | | **** | **** | *** |
| OVCAR10 | | | | | | | | | | **** | **** |
| A549 | | | | | | | | | | | **** |

| Figure 4b | MCF7 | MDAMB231 | PEO1 | SKOV3 | OVCA420 | OVCA429 | A2780 | HEYA8 | OVCAR10 | A549 | MKN28 |
|---|---|---|---|---|---|---|---|---|---|---|---|
| MCF10A | ns | ns | **** | ns | ns | ** | ns | ns | ns | ** | ns |
| MCF7 | | ns | **** | **** | ns | **** | ns | * | ns | **** | * |
| MDAMB231 | | | **** | *** | ns | **** | ns | ** | ns | **** | ns |
| PEO1 | | | | ** | **** | ** | **** | **** | **** | **** | **** |
| SKOV3 | | | | | ** | ns | **** | ** | ** | * | * |

| | | | | | | | | | | | |
|---|---|---|---|---|---|---|---|---|---|---|---|
| OVCA420 | | | | | | *** | ns | ns | ns | *** | ns |
| OVCA429 | | | | | | | **** | * | **** | ns | ** |
| A2780 | | | | | | | | ** | ns | **** | * |
| HEYA8 | | | | | | | | | * | ns | ns |
| OVCAR10 | | | | | | | | | | *** | ns |
| A549 | | | | | | | | | | | * |

**Supplementary Table 4 – Analysis of EphA receptor mRNA expression level.** Table reports the relative expression level of all EphA receptors relative to that of EphA2 (EphAx/EphA2) in cancer cell lines. With the exception for 4 cases (ration > 1) EphA2 is the most abundant receptor for ephrinA1.

| | EphA1 | EphA2 | EphA 3 | EphA 4 | EphA 5 | EphA 6 | EphA 7 | EphA 8 | EphA 10 |
|---|---|---|---|---|---|---|---|---|---|
| MCF7 | 0.51 | 1 | 0.14 | 10.02 | 0.14 | 0.15 | 0.59 | 0.27 | 0.36 |
| MCF10A | 0.01 | 1 | 0.00 | 0.02 | 0.00 | 0.00 | 0.00 | 0.00 | 0.00 |
| MDAMB231 | 0.01 | 1 | 0.01 | 0.14 | 0.01 | 0.01 | 0.01 | 0.02 | 0.02 |
| PEO1 | 0.53 | 1 | 0.02 | 0.05 | 0.02 | 0.02 | 0.02 | 0.07 | 0.06 |
| SKOV3 | 0.05 | 1 | 0.13 | 0.73 | 0.07 | 0.04 | 0.05 | 0.09 | 0.07 |
| OVCA420 | 0.22 | 1 | 0.07 | 0.14 | 0.05 | 0.04 | 0.04 | 0.09 | 0.16 |
| OVCA429 | 0.48 | 1 | 0.03 | 0.34 | 0.08 | 0.03 | 0.05 | 0.11 | 0.07 |
| A2780 | 0.08 | 1 | 4.66 | 0.07 | 0.06 | 0.06 | 3.02 | 0.16 | 0.08 |
| HEYA8 | 0.03 | 1 | 0.01 | 0.08 | 0.02 | 0.02 | 0.01 | 0.03 | 0.03 |
| OVCAR10 | 0.43 | 1 | 0.20 | 0.45 | 5.56 | 0.39 | 0.92 | 0.55 | 0.97 |
| A549 | 0.03 | 1 | 0.02 | 0.02 | 0.02 | 0.01 | 0.02 | 0.05 | 0.03 |
| MKN28 | 0.24 | 1 | 0.03 | 0.24 | 0.02 | 0.02 | 0.03 | 0.06 | 0.06 |

**Supplementary Table 5 –** List of gene positively correlating with EphA2 clustering.

| | EphA2 Radial Transport | | | Invasion potential | | |
|---|---|---|---|---|---|---|
| **Gene ID** | **p-value** | **FDR** | **Type of Correlation** | **p-value** | **FDR** | **Type of Correlation** |
| FOSL1 | 1.03E-06 | 7.59E-04 | + | 1.04E-06 | 1.17E-03 | + |
| SNRPG | 1.10E-06 | 7.59E-04 | + | 2.51E-03 | 1.07E-02 | + |
| PHLDA1 | 2.65E-06 | 9.18E-04 | + | 1.14E-04 | 4.46E-03 | + |
| TIMM23 | 3.17E-06 | 9.18E-04 | + | 4.70E-03 | 1.42E-02 | + |
| PLAU | 3.44E-06 | 9.18E-04 | + | 4.07E-03 | 1.33E-02 | + |
| MT1G | 4.36E-06 | 9.18E-04 | + | 5.04E-03 | 1.47E-02 | + |
| MT2A | 4.37E-06 | 9.18E-04 | + | 3.23E-03 | 1.20E-02 | + |
| CKAP5 | 6.05E-06 | 9.18E-04 | + | 3.64E-03 | 1.27E-02 | + |
| UBE2E3 | 6.89E-06 | 9.18E-04 | + | 5.29E-03 | 1.49E-02 | + |
| PTRF | 7.83E-06 | 9.18E-04 | + | 5.44E-04 | 6.28E-03 | + |
| ZCD1 | 8.22E-06 | 9.18E-04 | + | 6.66E-03 | 1.65E-02 | + |
| CAV2 | 8.34E-06 | 9.18E-04 | + | 2.52E-04 | 5.24E-03 | + |
| PHLDA1 | 8.87E-06 | 9.18E-04 | + | 4.06E-03 | 1.33E-02 | + |
| PI4KII | 9.42E-06 | 9.18E-04 | + | 1.25E-02 | 2.23E-02 | + |
| TAF1A | 1.09E-05 | 9.18E-04 | + | 4.33E-02 | 4.15E-02 | + |
| CAV1 | 1.12E-05 | 9.22E-04 | + | 1.32E-03 | 8.25E-03 | + |

| | | | | | | |
|---|---|---|---|---|---|---|
| GLS | 1.17E-05 | 9.32E-04 | + | 4.08E-03 | 1.33E-02 | + |
| TGFBR2 | 1.22E-05 | 9.32E-04 | + | 2.93E-04 | 5.52E-03 | + |
| MT1H | 1.39E-05 | 1.01E-03 | + | 1.58E-02 | 2.47E-02 | + |
| MAP4K4 | 1.74E-05 | 1.17E-03 | + | 3.42E-02 | 3.64E-02 | + |
| EPHA2 | 1.81E-05 | 1.20E-03 | + | 6.29E-05 | 3.88E-03 | + |
| ELK3 | 2.01E-05 | 1.26E-03 | + | 1.16E-03 | 7.97E-03 | + |
| BDNF | 2.10E-05 | 1.26E-03 | + | 1.53E-03 | 8.68E-03 | + |
| DLG7 | 2.19E-05 | 1.26E-03 | + | 1.66E-03 | 8.91E-03 | + |
| TMEM22 | 2.21E-05 | 1.26E-03 | + | 7.12E-03 | 1.70E-02 | + |
| CTSZ | 2.47E-05 | 1.26E-03 | + | 9.94E-03 | 1.96E-02 | + |
| PLAU | 2.77E-05 | 1.32E-03 | + | 3.23E-03 | 1.20E-02 | + |
| LHFP | 3.04E-05 | 1.33E-03 | + | 4.63E-04 | 6.07E-03 | + |
| EXT1 | 3.16E-05 | 1.37E-03 | + | 2.13E-02 | 2.86E-02 | + |
| CCDC99 | 3.26E-05 | 1.38E-03 | + | 6.18E-04 | 6.55E-03 | + |
| TNPO1 | 3.27E-05 | 1.38E-03 | + | 2.02E-04 | 4.88E-03 | + |
| XRCC5 | 4.11E-05 | 1.59E-03 | + | 5.22E-04 | 6.13E-03 | + |
| UPP1 | 4.13E-05 | 1.59E-03 | + | 2.60E-03 | 1.10E-02 | + |
| MT1E | 4.13E-05 | 1.59E-03 | + | 2.06E-03 | 9.86E-03 | + |
| NUP160 | 4.14E-05 | 1.59E-03 | + | 1.55E-02 | 2.44E-02 | + |
| EMP3 | 4.19E-05 | 1.59E-03 | + | 5.90E-04 | 6.41E-03 | + |
| PHLDA1 | 4.28E-05 | 1.60E-03 | + | 1.61E-03 | 8.82E-03 | + |
| AKAP12 | 4.38E-05 | 1.60E-03 | + | 1.51E-04 | 4.56E-03 | + |
| WDR79 | 4.66E-05 | 1.65E-03 | + | 1.75E-02 | 2.60E-02 | + |
| COTL1 | 4.72E-05 | 1.65E-03 | + | 2.17E-04 | 4.88E-03 | + |

| | | | | | | |
|---|---|---|---|---|---|---|
| ETV5 | 4.79E-05 | 1.65E-03 | + | 3.35E-03 | 1.21E-02 | + |
| HMGN4 | 4.85E-05 | 1.65E-03 | + | 1.12E-02 | 2.10E-02 | + |
| RTN4 | 5.06E-05 | 1.65E-03 | + | 4.35E-04 | 6.06E-03 | + |
| HBEGF | 5.10E-05 | 1.65E-03 | + | 1.94E-04 | 4.88E-03 | + |
| MYL6B | 5.14E-05 | 1.65E-03 | + | 9.50E-02 | 6.62E-02 | + |
| AKR1B1 | 5.19E-05 | 1.65E-03 | + | 1.84E-04 | 4.88E-03 | + |
| LDHB | 5.51E-05 | 1.68E-03 | + | 9.19E-03 | 1.91E-02 | + |
| Transcribed locus | 5.53E-05 | 1.68E-03 | + | 8.29E-03 | 1.83E-02 | + |
| PLAUR | 5.70E-05 | 1.70E-03 | + | 2.85E-02 | 3.32E-02 | + |
| TNPO1 | 5.88E-05 | 1.70E-03 | + | 8.78E-03 | 1.88E-02 | + |
| SH2B3 | 5.91E-05 | 1.70E-03 | + | 2.25E-03 | 1.02E-02 | + |
| ANXA1 | 5.94E-05 | 1.70E-03 | + | 3.07E-03 | 1.17E-02 | + |
| SLIT2 | 6.00E-05 | 1.70E-03 | + | 1.72E-02 | 2.58E-02 | + |
| EPS15 | 6.20E-05 | 1.72E-03 | + | 2.19E-04 | 4.88E-03 | - |
| AKAP2 | 6.44E-05 | 1.75E-03 | + | 3.02E-03 | 1.17E-02 | + |
| MCTP1 | 6.68E-05 | 1.77E-03 | + | 3.28E-03 | 1.20E-02 | + |
| WNT5A | 7.24E-05 | 1.86E-03 | + | 2.44E-03 | 1.07E-02 | + |
| GSTO1 | 7.39E-05 | 1.87E-03 | + | 3.44E-03 | 1.23E-02 | + |
| IRAK1 | 7.54E-05 | 1.90E-03 | + | 7.81E-02 | 5.87E-02 | + |
| CCT7 | 7.73E-05 | 1.92E-03 | + | 5.65E-03 | 1.54E-02 | + |
| CAV1 | 8.26E-05 | 1.97E-03 | + | 2.49E-03 | 1.07E-02 | + |
| TUBB6 | 8.30E-05 | 1.97E-03 | + | 3.14E-02 | 3.50E-02 | + |
| CDKN3 | 8.66E-05 | 2.00E-03 | + | 6.43E-03 | 1.61E-02 | + |
| GNG11 | 8.67E-05 | 2.00E-03 | + | 7.78E-04 | 6.82E-03 | + |

| | | | | | | |
|---|---|---|---|---|---|---|
| NUDC | 8.87E-05 | 2.01E-03 | + | 4.93E-03 | 1.46E-02 | + |
| PPARG | 8.88E-05 | 2.01E-03 | + | 9.91E-03 | 1.96E-02 | + |
| NOL8 | 8.88E-05 | 2.01E-03 | + | 9.04E-03 | 1.90E-02 | + |
| ISG15 | 9.13E-05 | 2.02E-03 | + | 7.17E-03 | 1.71E-02 | + |
| PSMC3 | 9.54E-05 | 2.09E-03 | + | 2.13E-02 | 2.86E-02 | + |
| PSMC4 | 9.76E-05 | 2.11E-03 | + | 6.27E-01 | 2.48E-01 | + |
| PNPLA6 | 9.77E-05 | 2.11E-03 | + | 3.67E-02 | 3.78E-02 | + |
